# Physiological levels of 3-hydroxykynurenine alter mitochondrial function and morphology in neuronal cells

**DOI:** 10.64898/2026.05.13.724856

**Authors:** Jordan Cassidy, Mary E W Collier, Flaviano Giorgini

**Affiliations:** Division of Genetics and Genome Biology, University of Leicester, University Road, Leicester, LE1 7RH, UK

**Keywords:** 3-hydroxykynurenine, mitochondria, kynurenine pathway, neurons, function, morphology, physiological, excitotoxicity, oxidative stress

## Abstract

Mitochondrial morphology and function are critical determinants of neuronal function and survival, with disruptions in mitochondrial dynamics often preceding the overt neuronal dysfunction seen in neurodegenerative diseases such as Alzheimer’s disease, Huntington’s disease and Parkinson’s disease. The kynurenine pathway accounts for 95% of dietary tryptophan catabolism and many of the metabolites are neuroactive, including redox-active 3-hydroxykynurenine (3-HK). 3-HK is present under normal physiological conditions in the central nervous system (CNS) and is elevated during inflammation. While supraphysiological levels of 3-HK have been associated with neurotoxicity, the effects of physiological concentrations on neuronal cells, and specifically their mitochondria, remain poorly understood. Here we assessed viability, ATP levels and redox status to determine cellular health and function in neuronal cells exposed to physiological levels of 3-HK, alongside confocal imaging and transcriptomic profiling, finding significant alterations in mitochondrial function and morphology. Interestingly, a biphasic influence of 3-HK on mitochondrial morphology was observed, with an elongated network as well as decreased surface area and volume being observed only at the lowest concentration of 3-HK, reflecting normal physiological levels. At the highest 3-HK concentration tested, reflecting an inflammatory situation, an increased number of mitochondria were present, accompanied by increased activation of caspase-3/7 and enhanced production of mitochondrial superoxide. These results highlight a previously unknown role for 3-HK in regulating mitochondrial function and structure, possibly through altered fission and fusion events, suggesting that subtle changes in kynurenine pathway metabolism may contribute to early mitochondrial dysfunction in neurological disease.

## Introduction

In mammals, the kynurenine pathway (KP) is the primary route for dietary tryptophan catabolism, accounting for 95% of its metabolism (Badawy, 2017). In the CNS, tryptophan is broken down by indoleamine 2,3-dioxygenase 1 (IDO-1) and IDO-2 to N-formyl-L-kynurenine (NFK) in the first step of the pathway (Ball *et al.,* 2014). NFK is hydrolysed to L-kynurenine (L-KYN), a process catalysed by arylformamidase (AFMID) (Dobrovolsky *et al.,* 2005), and L-KYN is then catabolised through two major branches: the terminal production of kynurenic acid (KYNA) by kynurenine aminotransferase (KAT) enzymes (Bellocchi *et al.,* 2009; Okuno *et al.,* 1991), or the intermediary 3-hydroxykynurenine (3-HK) by kynurenine-3-monooxygenase (KMO) in the central branch of the pathway (de Castro *et al.,* 1956; Makino *et al.,* 1954). 3-HK is further metabolised to 3-hydroxyanthranillic acid and quinolinic acid (QUIN) before the formation of nicotinamide adenine dinucleotide (NAD) occurs (Henderson, 1949; Mehler, 1956). Many of the metabolites of the kynurenine pathway have neuromodulatory roles, such as KYNA and QUIN which are primarily synthesised in astrocytes and microglia respectively and secreted to the extracellular milieu where they modulate neuronal glutamatergic signalling via N-methyl-D-aspartate receptor (NMDAR) activity (Guidetti *et al.,* 2007; Guillemin *et al.,* 2001; Heyes *et al.,* 1996; Smith *et al.,* 2007). 3-HK, the central metabolite of the KP has been shown to be present in the periphery of healthy individuals in the low nanomolar range (< 100 nM). However, increased 3-HK has been observed in neuropathologies, such as Huntington’s disease (HD) and Parkinson’s disease (PD) (Pearson & Reynolds, 1992; E. N. Wilson *et al.,* 2025).

The localisation of KMO to the outer mitochondrial membrane is evolutionarily conserved across a range of organisms including pigs, rats and yeast (Hirai *et al.,* 2010; Okamoto & Hayaishi, 1967; Zahedi *et al.,* 2006). In the brain, cellular localisation of KMO and subsequent production of 3-HK has been widely assumed to be “glio-centric”, where expression is primarily in microglial cells, with little to no expression found in astrocytes and uncertainty surrounding neuronal expression. However in recent years Castellano-Gonzalez *et al.,* (2019) found that 3-HK is readily produced by KMO in human primary neurons with Sathyasaikumar *et al.,* (2022) showing high activity of KMO in murine neurons (Castellano-Gonzalez *et al.,* 2019; Sathyasaikumar *et al.,* 2022). Nonetheless, an abundance of studies have shown that inflammatory stimulation of microglia increases KMO expression and activity hence neuronal access to 3-HK secreted by microglia in the brain following inflammatory stress is undisputed (Connor *et al.,* 2008; Esposito Soccoio *et al.,* 2026; Garrison *et al.,* 2018). It is becoming clear that neuronally produced 3-HK may also be present during infection and neuroinflammation where infiltrating macrophages with high KMO expression enter the brain (Alberati-Giani *et al.,* 1996); hence, neurons are exposed to 3-HK from a range of sources that fluctuate following inflammation.

Previous studies have largely focused on the neurotoxic effects of supraphysiological levels of 3-HK, particularly in relation to oxidative damage and cell death. Cell-based studies use 3-HK in the high micromolar/low millimolar range despite its presence in the human periphery at a low nanomolar concentration. Although 3-HK is elevated in diseases such as HD, PD and depression, neurons are routinely exposed to lower concentrations of 3-HK in normal conditions and a redox modulatory and non-neurotoxic role has been suggested (Colín-González *et al.,* 2013, 2014; Goshima *et al.,* 1986; Leipnitz *et al.,* 2007; Luthra & Balasubramanian, 1992). Despite this, the role of lower concentrations of 3-HK in neurons, and in particular its effect on mitochondrial function following production by mitochondrial enzyme KMO, remain poorly understood (Hirai *et al.,* 2010).

Mitochondrial morphology is a critical determinant of neuronal function, reflecting the dynamic balance between mitochondrial fission and fusion processes that support cellular energy demands and reduction-oxidation (redox) regulation. A combination of fission and fusion events, and cytoskeletal interactions regulate mitochondrial morphology (Palmer *et al.,* 2011). In cells with high energy demands, mitochondria fuse to form elongated networks, efficient at producing adenosine triphosphate (ATP). However, following cellular stress, fission is upregulated to isolate damaged mitochondria for mitophagy (Srinivasan *et al.,* 2017; Vásquez-Trincado *et al.,* 2016). The actin cytoskeleton oversees tight control of mitochondrial fission at endoplasmic reticulum (ER)-mitochondria contact sites through actin polymerisation constricting the mitochondria, before factors such as dynamin-related protein 1 (Drp1) are recruited to complete the scission (Friedman *et al.,* 2011; Hatch *et al.,* 2014; Ji *et al.,* 2015).

Disruptions to mitochondrial dynamics (fission, fusion and transport) have been increasingly implicated in the pathogenesis of neurodegenerative disorders, including Alzheimer’s disease (AD) (Devi *et al.,* 2006; Saraiva *et al.,* 1985), HD (Liu *et al.,* 2022; Shirendeb *et al.,* 2012) and PD (C. Chen *et al.,* 2023; Schapira *et al.,* 1990), where altered mitochondrial morphology often precedes neuronal dysfunction. This is likely due to mitochondria having a critical multi-faceted role in maintaining neuronal function (Tong *et al.,* 2026). Neurons are cells with high energy requirements and require mitochondrial ATP produced through oxidative phosphorylation to maintain membrane potential and synaptic transmission (Rumpf *et al.,* 2023). As neurons can have long axons, mitochondria are transported towards the synapses and growth cones for enhanced local ATP production (Hollenbeck & Saxton, 2005). Mitochondrial metabolism supplies metabolites used in neurotransmitter synthesis, redox balance and lipid metabolism (Frezza, 2017). During neuronal activity, calcium influx occurs and intracellular calcium is regulated by mitochondrial calcium uniporters to prevent the toxic accumulation of calcium which could lead to cell death (Beccano-Kelly *et al.,* 2023; Rizzuto *et al.,* 2012). Moreover, mitochondria directly control pathways which affect neuronal survival and influence the removal of damaged neurons such as the initiation of apoptosis through cytochrome c release (Kalkavan & Green, 2018; Tait & Green, 2010).

In this study, we investigated the effects of physiological concentrations of 3-HK on mitochondrial function and morphology in a neuronal cell model. We mimicked the *in vivo* environment where neighbouring microglial cells release 3-HK to neurons. Our neuronal cell model had almost no detectable *KMO* expression, hence our paradigm mimics a “glio-centric” exposure of neurons to 3-HK. We used physiologically relevant concentrations of 3-HK to model the *in vivo* environment, where elevated 3-HK is both representative of healthy and inflammatory conditions. As inflammation is a physiological response, we refer to our concentrations as physiological throughout in both ranges that would in theory encompass healthy and diseased states. We chose 100 nM (0.1 µM) to reflect a healthy, non-inflamed environment whilst 1 µM and 10 µM were chosen to mimic a progressively inflamed state.

## Materials & Methods

### Mammalian cell culture

The human neuroblastoma cell line SH-SY5Y (ECACC 94030304) was cultured in DMEM containing 10% (v/v) foetal bovine serum (FBS) at 37 °C under 5% CO_2_. Prior to experimental use, media was exchanged to DMEM containing 5% (v/v) FBS and cells were incubated for 24 hours at 37 °C. For experiments, cells were seeded into 96-well plates (2 x 10^4^) or T75 flasks (1 × 10^6^ cells) in DMEM containing 2% (v/v) FBS and incubated for 24 hours at 37 °C. Cells were washed twice with phosphate buffered saline (PBS), pH 7.4, before the addition of serum-free media (SFM), containing recombinant nerve growth factor (NGF) (50 ng/mL) (Thermo Fisher Scientific, Loughborough, UK), alongside the addition of 3-hydroxy-DL-kynurenine (Sigma-Aldrich) for 24 hours. Unless otherwise stated, a stock solution of 5 mM 3-HK was prepared fresh on the day of the experiment by dissolving directly into SFM. 3-HK was protected from light in all instances. Cells were treated with 0.1, 1 or 10 µM 3-HK unless otherwise stated.

### Differentiation of human neuroblastoma SH-SY5Y cells

Cells were seeded into 96-well plates at a density of 5 x 10^3^ in DMEM supplemented with 5% FBS and incubated at 37 °C for 24 hours. Differentiation occurred following the protocol described by Encinas *et al.,* (2000) with minor modifications. Cells were treated with 10 µM of all-trans retinoic acid (RA) (Sigma) in DMEM containing 5% FBS and incubated for 48 hours. Cells were treated with 10 µM RA in DMEM containing 2% FBS and incubated for a further 48 hours. Cells were then placed in SFM supplemented with 10 µM RA and 12.5 ng/mL brain-derived neurotrophic factor (BDNF) (Cambridge Biosciences, Cambridge, UK). The RA/BDNF-supplemented SFM was then replaced every other day up to day 10. On day 11, cells were washed twice with 200 µL of PBS, before the addition of 100 µL SFM, containing NGF (50 ng/mL) alongside the addition of 3-HK for 24 hours.

### SDS-PAGE and Immunoblot analysis

20 µg SH-SY5Y cell lysates were separated on a 10-20% (w/v) SDS-PAGE pre-cast Novex™ Tris-Glycine Mini Protein Gel (Invitrogen, XP10202BOX), then transferred onto nitrocellulose membranes and blocked with Tris-buffered saline Tween20 (TBST) (50 mM Tris-HCl, pH 8, 150 mM NaCl, 1% (v/v) Tween-20) containing 5% (w/v) low fat milk for 1 hour. Membranes were incubated overnight at 4 °C with the following primary antibodies diluted in TBST-milk; rabbit monoclonal Anti-Enolase-2 (Bio-Techne; clone E2H9X, 1:1000 dilution) and rabbit anti-MAP2 (Proteintech; 17490-1-AP, 1:5000). Membranes were washed three times for 5 minutes each with TBST and then incubated for 1 hour at room temperature with an anti-rabbitHRP conjugated secondary antibody (Vector Labs, Kirtlington, UK; 1:5000). Membranes were washed four times with TBST and developed with Westar Supernova ECL detection reagent (Cyanagen, Bologna, Italy) and imaged using a G: BOX Chemi XX9 gel imaging system (Syngene, Cambridge, UK).

### Crystal violet cell viability assay

Cell viability of SH-SY5Y cells was determined using the DNA intercalating dye, Crystal Violet (Thermo Scientific, 447570500) at 0.125% (w/v) in 20% methanol solution made using distilled water. 24 hours post-treatment of cells with 3-HK (0 – 500 µM), the media was discarded from 96-well plates, and the cells were washed with 100 µL of PBS. Crystal violet solution was added to the plates at 50 µL/well and incubated at room temperature (RT) for 10 minutes before being washed using distilled water until no purple coloration of the water was observed. Open plates were left to dry before solubilisation of the stain using 70% ethanol at 100 µL/well. Plates were placed upon an orbital microplate shaker at 650 rpm for 10 minutes before absorbance at 570nm was recorded using a FLUOstar Omega microplate reader.

### Caspase-3/7 Assay

To determine caspase-3/7 activation following 3-HK inclusion, the luminescent-based Caspase-Glo® 3/7 Assay (Promega, G8090) was used in accordance with the manufacturers’ instructions. SH-SY5Y cells were seeded into white 96-well plates in 100 µL of DMEM containing 2% FBS and incubated at 37°C, 5% CO_2_ for 24 hours. A full media exchange was then carried out and 3-HK was added to each well in 100 µL of SFM supplemented with 50 ng/mL NGF. Cells were incubated at 37°C for 24 hours. Prior to the addition of Caspase-Glo® reagent, plates were allowed to equilibrate to RT for 30 minutes Media was then discarded, and wells were washed once using 100 µL pre-warmed SFM. 25 µL SFM was added to each well followed by an equal volume of Caspase-Glo® reagent. Plates were gently shaken for 2 minutes and then the plate was incubated for 30 minutes at RT for the luminescent signal to stabilize. Luminescence was recorded using a FLUOstar Omega microplate reader.

### MitoSOX staining

SH-SY5Y cells were seeded at 1 x 10^4^ cells/chamber in a CELLview^TM^ 35 mm cell culture dish (Greiner, 627870) in DMEM with 2% FBS and incubated at 37°C for 24 hours. A full media exchange was then carried out and 3-HK was added to each chamber in 500 µL of SFM supplemented with NGF. All chambers contained a final concentration of 0.02% dimethyl sulfoxide (DMSO), including a 3-HK naïve vehicle control chamber. Cells were incubated with 3-HK at 37°C for 24 hours before the media was refreshed with 250 µL of pre-warmed SFM including a final concentration of 4 µM MitoSOX Red (Invitrogen, M36008) and 2.5 µM of Hoechst 33342 (Abcam, ab228551). Cells were incubated at 37°C, 5% CO_2_ in the dark for 10 minutes before the media was discarded and cells were rinsed using fresh SFM and placed in 500 µL of FluoroBrite™ DMEM (Gibco, A1896701). Imaging was performed immediately on the live cells in a viewing chamber maintained at 37°C, 5% CO_2_. Imaging of stained samples took place using the VisiTech HAWK system; an Infinity 3 confocal microscope (VisiTech International). Acquisition and image capture was performed using the VoxCell Scan acquisition software, version 2.0 (VisiTech International). Images were captured at 60X magnification using lasers at 405 nm (DAPI) and 561 nm (Rhodamine). In both channels, zeta-stacks were captured at step intervals of 0.2 µm. Blinded image analysis was performed using FIJI on maximal projection images compiled from each zeta stack (Schindelin *et al.,* 2012). Fluorescence intensity was measured as the integrated density (AU) in the rhodamine channel and divided by the number of nuclei per image present in the DAPI channel.

### Metabolic activity assay

SH-SY5Y cells were seeded into black 96-well plates at 2 x 10^4^ cells/well in 100 µL of DMEM containing 2% FBS and incubated at 37°C for 24 hours. A full media exchange was then carried out and 3-HK was added to each well in 100 µL of SFM supplemented with NGF and incubated at 37°C for 24 hours. The media was then discarded and cells were washed once using pre-warmed SFM. 100 µL of the redox-sensitive dye resazurin (MedChemExpress, Cat. No. HY-118540) (44 µM) in SFM was added to each well and incubated for 2 hours at 37°C, 5% CO_2._ Fluorescence at 544nm_EX_/690nm_EM_ was recorded using a FLUOstar Omega microplate reader.

### ATP measurements

To determine ATP production following incubation of cells with 3-HK, the CellTiter-Glo® 2.0 Cell Viability Assay (Promega, G9241) was used in accordance with manufacturers’ instructions. SH-SY5Y cells were seeded into white 96-well plates at 2 x 10^4^ cells/well in 100 µL of DMEM with 2% FBS and incubated at 37°C for 24 hours. A full media exchange was then carried out and 3-HK was added to each well in 100 µL of SFM supplemented with NGF and incubated at 37°C for 24 hours. Prior to the addition of CellTiter-Glo® reagent, plates were allowed to equilibrate to RT for 30 minutes. Media was then discarded and cells were washed once using pre-warmed SFM. 25 µL SFM was added to each well followed by an equal volume of CellTiter-Glo® reagent. Plates were gently shaken for 2 minutes and then the luminescent signal was allowed to stabilise for 10 minutes at RT. Luminescence was recorded using a FLUOstar Omega microplate reader.

### Seahorse Mito Stress Assay

The Seahorse XFp Extracellular Flux Analyzer (Agilent Technologies, Santa Clara, CA, USA) was used to generate the bioenergetic profiles of SH-SY5Y cells upon treatment with 10 µM 3-HK. Live-cell analyses of oxygen consumption rate (OCR) was measured using the Mito Stress test (Agilent). Cells were cultured on a Seahorse XFp cell culture micro-plate (Seahorse Biosciences; Agilent Technologies) at a density of 2.5 × 10^4^ cells/well and grown overnight in 100 µl DMEM and 2% FBS. The companion sensor cartridge was hydrated with calibration solution (Seahorse Biosciences; Agilent Technologies) for 24 hours in a non-CO2 incubator at 37 °C. The following day, one hour prior to the assay, the cell culture media was replaced with 200 µl of pre-warmed XF assay medium (Seahorse Biosciences; Agilent Technologies) supplemented with 1 mM pyruvate, 2 mM GlutaMAX and 10 mM glucose. The plate was then placed in a 37°C, non-CO2 incubator for 1 hour until assay commencement.

The Seahorse bioanalyzer XFp was run on the Mito Stress program pre-saved to the Wave software as per the manufacturer’s guidance (Seahorse Biosciences). After baseline measurements for the oxygen consumption ratio, cells were sequentially challenged with injections of Mito Stress drugs prepared following the manufacturer’s instructions. The final concentrations used for each drug were 7.5 μM oligomycin (ATP synthase inhibitor), 4 μM carbonyl cyanide-p-trifluoromethoxyphenylhydrazone (FCCP; mitochondrial respiration uncoupler), and 1 μM rotenone/antimycin (complex I and III inhibitors). Data was exported to Seahorse Analytics (Agilent) and extracted using Wave software. Immediately following program completion, cells were directly lysed in 30 µL radioimmunoprecipitation assay (RIPA) buffer (50 mM Tris-HCl, pH7.4, 150 mM NaCl, 0.5% (w/v) sodium deoxycholate, 0.1% (w/v) sodium dodecyl sulphate (SDS), 1% (v/v) Triton X-100, 1% (v/v) Halt protease inhibitor cocktail (100X)) (Themo Scientific, 87786) for quantification of total protein content using the detergent compatible (DC) Protein Assay (Bio-Rad), following manufacturer’s instructions. OCR measurements were normalised to total protein content per well (pmol O_2_/minute/µg protein).

### Mitochondrial staining with Mitoview^TM^ Green Dye

SH-SY5Y cells were seeded at 1 x 10^4^ cells/chamber in a CELLview^TM^ cell culture dish in 500 µL of DMEM with 2% FBS and incubated at 37°C for 24 hours. A full media exchange was then carried out and 3-HK (0-10 µM) was added to each chamber in 500 µL of SFM supplemented with NGF. All chambers contained a final concentration of 0.02% DMSO. Cells were incubated with 3-HK treatments at 37°C for 24 hours before media was refreshed as 250 µL pre-warmed SFM including a final concentration of 60 nM MitoView^TM^ Green (Biotium, 70054-T), a mitochondrial membrane potential independent dye. Cells were incubated at 37°C in the dark for 20 minutes before the media was discarded and cells were rinsed using fresh SFM before being replaced for a final time with 500 µL of FluoroBrite™ DMEM (Gibco, A1896701). Imaging was performed immediately on the live cells in a viewing chamber maintained at 37°C, 5% CO_2_ using the VisiTech HAWK system; an Infinity 3 confocal microscope (VisiTech International).

Acquisition and image capture was performed using VoxCell Scan acquisition software, version 2.0 (VisiTech International). Images were captured at 60X magnification using the laser channel 488 nm (GFP). Zeta-stacks were captured at step intervals of 0.2 µm.

### Analysis of MitoView ^TM^ staining for mitochondrial morphology

Blinded image analysis of MitoView^TM^ stained cells, focusing on mitochondrial network morphology and individual mitochondria shape and number, was performed using FIJI (Schindeln *et al.,* 2012) following an adjusted method based on that done by Chaudhry *et al.,* (2019). To begin, image stacks were blinded and then underwent deconvolution using the Deconvolution Lab 2 FIJI plugin (Sage *et al.,* 2017). Each image in the stack then underwent processing using the following commands: ‘subtract background’ (radius = 1 μm) to remove background noise; ‘sigma filter plus’ (radius = 0.1 μm, 2.0 sigma) to reduce noise and smooth object signal while preserving edges; ‘enhance local contrast’ (block size = 108, histogram = 256, slope = 1.25) to enhance dim areas while minimizing noise amplification; ‘gamma correction’ (value = 1) to correct any remaining dim areas; ‘8-bit’ to enable further processing using greyscale; ‘Auto Local Threshold’ (method=Bernsen radius=15) to create a threshold based per pixel; ‘Despeckle’ to reduce noise; and ‘remove outliers’ (radius = 0.2 μm, maximum = 50) to further remove residual noise. Image stacks were then manually inspected for quality and individual cells were traced using the polygon tool. Tracings were duplicated twice for cellular analysis using one of two macros, detailed as follows: both macros started with the commands ‘Clear outside’ to remove noise outside the area of interest and ‘3D objects counter’ to initialise analysis of each image in the stack. To derive the number of mitochondria per cell, volume, surface area and sphericity the command ‘particle analysis 3D’ was enacted. Otherwise, to obtain a skeletal outline of the mitochondrial network for determination of the number of branches, branch length and branch junctions the commands ‘Skeletonise (2D/3D)’ and ‘Analyse skeleton (2D/3D)’ were performed. Data was then entered into a spreadsheet for all 3 independent experiments before image stacks were unblinded and analysed.

### F-actin staining

SH-SY5Y cells were seeded at 1 x 10^4^ cells/chamber in a CELLview^TM^ 35 mm cell culture dish in DMEM with 2% FBS and incubated at 37°C for 24 hours. A full media exchange was then carried out and 3-HK was added to each chamber in 500 µL of SFM supplemented with NGF. All chambers contained a final concentration of 0.02% DMSO, including a 3-HK naïve vehicle control chamber. Cells were incubated with 3-HK at 37°C for 24 hours before the media was discarded and cells were washed thrice with 500 µL of ice-cold 0.1% bovine serum albumin (BSA) in PBS before 200 µL of 4% paraformaldehyde (PFA) solution was added per well. Cells were incubated in fixative at RT for 20 minutes before a further three washes with 0.1% BSA occurred. 200 µL of staining solution containing 165 nM Alexa Fluor 594 phalloidin (Invitrogen, A12381) was added to each chamber and incubated at RT in darkness for 20 minutes. Staining solution was discarded and cells were washed thrice more with 0.1% BSA before 500 µL of fresh 0.1% BSA was added to each chamber and dishes were kept at 4°C until imaging occurred later that same day. Imaging of stained samples took place using the VisiTech HAWK system; an Infinity 3 confocal microscope (VisiTech International). Acquisition and image capture was performed using the VoxCell Scan acquisition software, version 2.0 (VisiTech International). Images were captured at 100X magnification using a laser at 561 nm (Rhodamine) and zeta-stacks were captured at step intervals of 0.2 µm. Blinded image analysis was performed using FIJI on maximal projection images compiled from each zeta stack (Schindelin *et al.,* 2012). Fluorescence intensity was measured as the integrated density (AU) in the rhodamine channel.

### RNA extraction and mRNA sequencing

SH-SY5Y cells were seeded at 1 x 10^6^ cells in T75 flasks, adapted to SFM and treated with 3-HK for 24 hours. The final concentration of DMSO was 0.02% in each sample. RNA was extracted from cells using the TRIzol Reagent according to the manufacturer’s guidelines (Invitrogen) and Phasemaker tubes. The RNA pellet was resuspended in 40 µL of RNAse-free water and incubated on a 55 °C heat block for 15 minutes for solubilisation, before performing RNA quantification, quality assessment and downstream analysis. The quality and concentration of the extracted RNA was determined using the nanophotometer (IMPLEN, P300) and Agilent RNA 6000 Nano kit on an Agilent 2100 Bioanalyzer system. Samples with an OD260/230 ≥ 1.8 and a concentration > 10 ng/µL proceeded to be sent to Novogene Cambridge Genomic Centre (Novogene Company Ltd, Cambridge, UK) for further quality assessment (RNA integrity number ≥ 9), mRNA library preparation (poly A enrichment) and sequencing using Illumina PE150, generating at least 20M reads per sample.

### Quantitative PCR for KP enzyme expression

Contaminating genomic DNA was removed from the RNA samples (4 µg) using the TURBO DNA-free™ Kit (Ambion, Life technologies, Cat. No. AM1907) according to the manufacturer’s instructions. cDNA synthesis was then performed using QuantiTect Reverse Transcription Kit (Cat. No. 205311). 2 µL of gDNA wipeout buffer was added to each 12 µL reaction and incubated for 2 minutes at 42°C. 4 µL of reverse transcriptase buffer and 1 µL reverse transcriptase primer mix were added to all tubes. 1 µL of reverse transcriptase enzyme was then added to 3 of the tubes and 1 µL RNase-free water was added to the final tube as a reverse transcriptase null control. Reactions were incubated for 15 minutes at 42°C for cDNA synthesis, then 3 minutes at 95°C to inactivate the reverse transcriptase enzyme. Samples were then stored at -20°C until use.

All primer sequences are shown in Supplementary Table 1. Gene expression was quantified using the QuantiTect SYBR Green PCR Kit (Cat. No. 204143) according to manufacturer’s instructions. The following was added to each well: 1 μL cDNA, 0.5 μM of each forward and reverse primer, 7 μL SYBER Green Master Mix, and RNase free water to a final volume of 10 μL per well. Three biological replicates were dispensed into triplicate wells and the run was performed in a Roche Light Cycler® 480 using the LightCycler software, version 1.5.0, using the following cycling conditions: 1) initial denaturation: 95 °C for 15 minutes; 2) denaturation: 95 °C for 15 seconds; 3) annealing: 60 °C for 1 minute; 4) extension: 72 °C for 30 seconds (data acquisition occurred at this step). Steps 2-4 were repeated for 40 cycles.

### Bioinformatic Analyses

All bioinformatics were performed using the ALICE High Performance Computing facility at the University of Leicester. Analysis was performed using R, version 4.2.2 (R Core Team, 2023), and Rstudio, version 2023.9.0.463 (Posit Team, 2023).

Read quality was evaluated using FastQC 0.12.9 (Andrews, 2010) and summarised using MultiQC 1.12 (Ewels *et al.,* 2016). Reads were then aligned, using HISAT2 version 2.2.1 (Kim *et al.,* 2019), to the human reference genome from UCSC (hg38, http://igenomes.illumina.com.s3-website-us-east-1.amazonaws.com/Homo_sapiens/UCSC/hg38/Homo_sapiens_UCSC_hg38.tar.gz), downloaded from Illumina igenomes. Transcript abundance was counted using StringTie 2.2.1 (Pertea *et al.,* 2015) and the assembled transcriptomes were quantified using the prepDE.py3 script provided by the StringTie developer to generate transcript counts for downstream analysis. Differential transcript expression (DTE) between experimental conditions was evaluated using the DESeq2 R package, version 1.36.0 (Love *et al.,* 2014) using R version 4.2.2 (R Core Team, 2022). After filtering of transcripts, those with low read counts (< 10) were discarded and transcript counts were normalised by library and condition. Following the Benjamini-Hochberg method (Benjamini & Hochberg, 1995), P-values were adjusted to control the false discovery rate. The dataset list obtained was filtered selecting genes with an adjusted P-value ≤ 0.05 and a minimum of 1.5 log fold-change (logFC). The scripts used are available at: https://github.com/jordanxcassidy.

### Gene ontology functional classification using PANTHER

Differentially expressed transcripts (DETs) between all experimental conditions were combined and evaluated using PANTHER classification software, version 19.0 (Mi *et al.,* 2013). Gene ontology enrichment and functional classification of DETs was performed through uploading the list of significant DETs, gained from DESeq2 and selection of *Homo sapiens* as the organism. The false discovery rate was controlled using the Benjamini-Hochberg correction (Benjamini & Hochberg, 1995).

### Statistical Analysis

Statistical analysis was performed using GraphPad Prism software version 9.4.0 (GraphPad Inc. San Diego, CA). Specific details of performed tests are noted in respective figure legends. Statistical significance was deemed when the P-value was < 0.05.

## Results

### Exogenous 3-HK induces cellular toxicity at high concentrations in non-differentiated but not differentiated SH-SY5Y cells

We first investigated whether non-differentiated or differentiated cells would be most suitable for our study as SH-SY5Y cells can be differentiated into a more mature neuronal phenotype (Encinas *et al.,* 2000). Treatment of cells with RA and BDNF resulted in the differentiation of cells into a neuronal phenotype as shown by Encinas *et al.,* (2000) with extended neurites observed compared to non-differentiated cells and the upregulation of neuron specific enolase and MAP2 protein expression (Supp. Figure 1A-C). qPCR analysis of KP genes expressed in neuroblastoma cells revealed that *KMO*, the enzyme responsible for 3-HK synthesis, was not detected in differentiated cells and expressed at levels close to the limit of detection (determined by the “no reverse transcriptase” control) in non-differentiated cells. Of the 11 KP enzymes examined, Ct values indicated that four had detectable expression levels in both undifferentiated and differentiated SH-SY5Y cells: *AFMID*, *KYAT1*, *AADAT* and *KYAT3*. Detection of housekeeping genes *IPO8* and *Rplp0* was also confirmed (Supp. Table 2). From published studies, 3-HK up to and including 10 µM was deemed a physiologically relevant concentration (to include healthy and inflammatory ranges), while greater than 10 µM was taken as supraphysiological. For our initial viability studies, we assessed the effects of both physiological and supraphysiological levels of 3-HK. The crystal violet cell viability assay showed no changes in the viability of differentiated cells at any of the 3-HK concentrations tested when compared to the untreated control cells (Figure 1A). As we observed no changes in the viability of differentiated cells even at the highest concentration of 3-HK (250 µM), we chose to pursue only the non-differentiated cell line as a model. No phenotypic changes following treatment of non-differentiated cells with 3-HK was observed up to 50 µM, but cell detachment and diminished processes were present at 100 µM 3-HK (Supp. Figure 1D). Moreover, a significant decline in cell viability was observed in non-differentiated cells treated with ≥ 100 µM 3-HK (Figure 1A). Despite no significant impact on viability being observed following 3-HK treatments below 100 µM 3-HK, increased activity of caspase-3/7 was observed when non-differentiated cells were treated with ≥ 1 µM 3-HK relative to the untreated control as well as the 0.1 µM 3-HK treatment (Figure 1B). As we were most interested in recapitulating the *in vivo* environment, we chose to move forward with physiologically relevant concentrations of 3-HK only.

**Figure 1:**
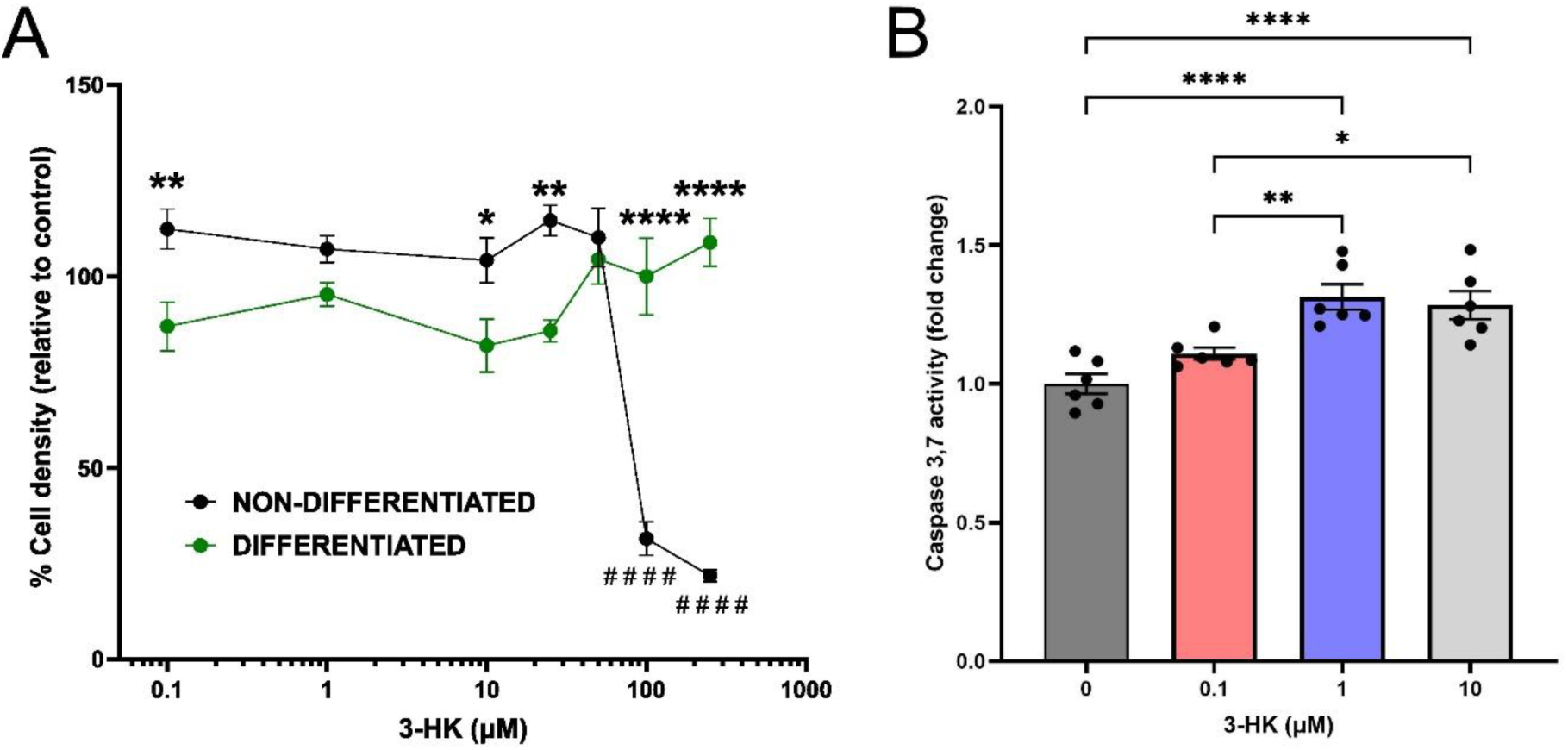
Exogenous 3-HK induces cellular toxicity at high concentrations in non-differentiated but not differentiated SH-SY5Y cells. (A) Crystal violet assay: Following 24-hours incubation with 0 - 250 µM 3-HK, the percentage cell density of non-differentiated SH-SY5Y cells and differentiated SH-SY5Y cells was compared. ****P < 0.0001. **P < 0.01. *P < 0.05. The percentage cell density of non-differentiated SH-SY5Y cells was compared, relative to the untreated control. ####P < 0.0001. N = 8. Two-way AVOVA with Šídák’s multiple comparisons test. Error bars indicate ± SEM. (B) Caspase-3/7 activity assay: Following addition of Caspase-Glo®, luminescence values were recorded using a FLUOstar Omega plate reader (BMG LabTech) and used to calculate caspase-3/7 activation. Data are shown as fold change relative to the drug-naïve control. N = 6. Data points represent a single well and error bars indicate ± SEM. One-way ANOVA with Tukey’s multiple comparison test. ****P < 0.0001. **P < 0.01. *P < 0.05.

### Physiological levels of exogenous 3-HK increase cellular redox activity but not ATP production

The observed increase in caspase-3/7 activation indicated cellular stress induced by 3-HK, which led us to further evaluate the effects of 3-HK on cellular mechanisms related to cell viability. As cell viability can be altered by metabolic and oxidative stress (Annunziato *et al.,* 2003; Zhou *et al.,* 2019), we assessed both metabolic activity and redox indicators within the 3-HK treated SH-SY5Y cells. Cellular metabolic activity was assessed using two distinct metrics: cellular redox activity and ATP production. Redox activity was measured through quantification of the reduction of the redox sensitive dye resazurin to resorufin which showed a significant increase in redox activity following treatment with 1 µM 3-HK, compared with the 0.1 µM 3-HK condition (Figure 2A). However, no change in cellular ATP production was observed when cells were treated with a physiologically relevant range of 3-HK concentrations (Figure 2B).

**Figure 2:**
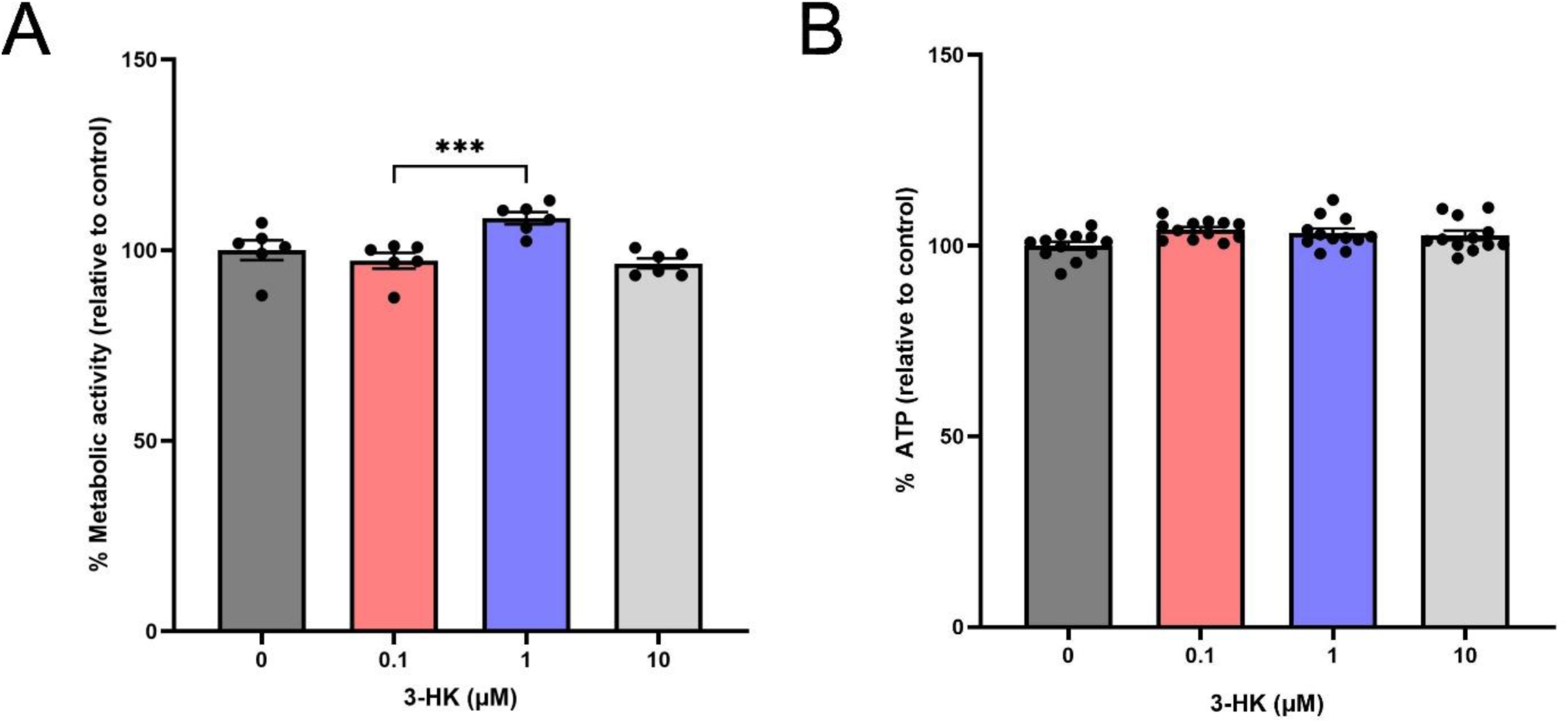
Physiologically relevant levels of exogenous 3-HK increased cellular metabolic activity but not ATP production. *(A)* Reduction of resazurin to resorufin by SH-SY5Y cells was significantly increased following 24 hours exposure to 1 μM 3-HK, compared with the 0.1 μM 3-HK condition. Following addition of resazurin, fluorescence at 544nm_EX_/690nm_EM_ was recorded using a FLUOstar Omega microplate reader (BMG LabTech). Fluorescence values were used to calculate the percentage (%) of resorufin produced, relative to the drug-naïve control. This was used as an indirect indication of mitochondrial metabolic activity (%). N = 6. Data points represent a single well and error bars indicate ± SEM. One-way ANOVA with Tukey’s multiple comparisons test. ***P < 0.001. *(B)* ATP production by SH-SY5Y cells was unchanged following 24 hours exposure to 3-HK. Following the addition of CellTiter-Glo®, luminescence values were recorded using a FLUOstar Omega microplate reader (BMG LabTech) and used to calculate the percentage (%) of ATP produced relative to the drug-naïve control. N = 12. Data points represent a single well and error bars indicate ± SEM. Assays were performed in duplicate. One-way ANOVA with Tukey’s multiple comparisons test.

### Exogenous 3-HK increases mitochondrial superoxide production in neuroblastoma cells

Redox activity and ATP production are indicators of mitochondrial health but are not direct measurements of mitochondrial activity due to the contributions of cytosolic enzymes and glycolysis (Zheng, 2012). However, as endogenous 3-HK is produced by KMO in the outer mitochondrial membrane and released to the immediate surrounding cytosol, we chose to directly evaluate mitochondrial function within 3-HK treated SH-SY5Y cells. Mitochondrial functionality was therefore assessed by measuring mitochondrial superoxide production. Confocal microscopy images of cells treated with increasing concentrations of 3-HK and then incubated with MitoSOX indicated that mitochondrial superoxide was present in all conditions and was localised to the perinuclear region of cells (Figure 3A). Superoxide levels were slightly increased in cells treated with 0.1-1 µM of 3-HK compared to untreated controls, but significantly increased levels of mitochondrial superoxide were observed in SH-SY5Y cells following treatment with 10 µM 3-HK when compared with the 3-HK-naïve control (Figure 3B).

**Figure 3:**
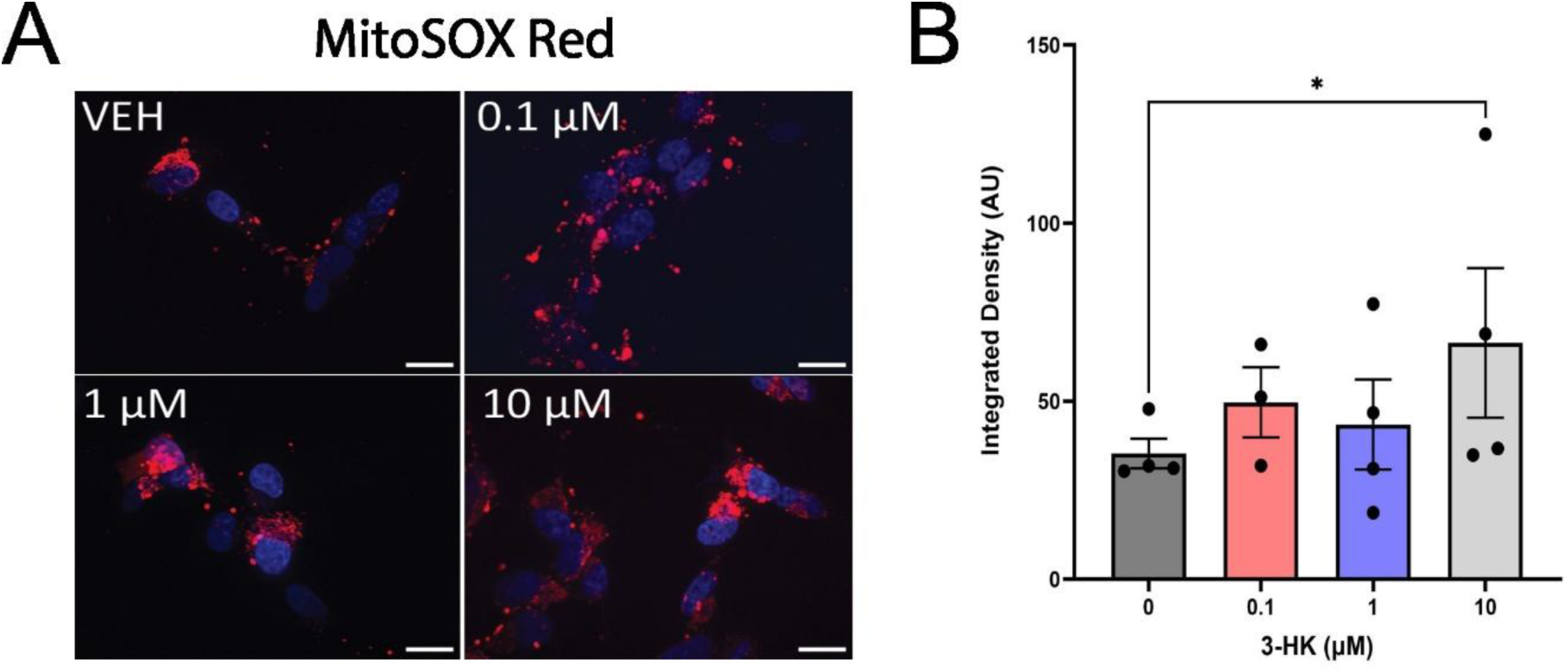
Mitochondrial superoxide production was increased in SH-SY5Y cells following 24 hours exposure to 10 μM 3-HK. Fluorescence intensity (AU) of MitoSOX Red staining was measured by confocal microscopy to quantify production of mitochondrial superoxide in SH-SY5Y cells treated with 0.1 – 10 μM 3-HK. (A) Representative confocal microscopy images for each condition. Red = MitoSox Red. Blue = nuclei (Hoechst 33342). Magnification x60. Scale bar = 20 μm. *VEH = vehicle control. (B)* Quantification of confocal images. N = 4. Points represent the average value per replicate from ≥ 3 independent assays. Two-way ANOVA with Dunnett’s multiple comparisons test. Error bars indicate ± SEM. *P < 0.05.

Next, mitochondrial respiration was assessed following treatment of cells with 10 µM 3-HK using the MitoStress test on the Seahorse analyser (Agilent Biosciences). This determines real-time metabolic activity of cells through measuring their OCR as they are exposed to specific inhibitors of the mitochondrial electron transport chain. However, no significant alterations in OCR, basal respiration, ATP-linked respiration, non-mitochondrial respiration or coupling efficiency were observed (Supp. Figure 2.), indicating that mitochondrial respiration was not affected by 3-HK.

### 3-HK alters mitochondrial number and surface area to volume ratio in a concentration dependent manner

Due to the altered metabolic properties of the neuroblastoma cells observed when treated with physiologically relevant levels of 3-HK, and the known link between mitochondrial functionality and morphology (H. Chen *et al.,* 2005; Singh *et al.,* 2025; Yu *et al.,* 2006), we next investigated mitochondrial size, shape and number in response to 3-HK. Mitochondrial morphology in MitoView^TM^-stained cells was examined in detail by ImageJ analysis of confocal microscopy images. We observed that samples without 3-HK treatment had both branching, tubular mitochondria connected in networks as well as distinct punctate mitochondria, indicative of a balance between both fission and fusion events. Interestingly, samples treated with 0.1 μM 3-HK had an almost entirely connected mitochondrial network, lacking any distinct punctate or rod structures, whereas samples treated with 1 μM or 10 μM 3-HK displayed a mitochondrial network which became increasingly fragmented, lacking branching as 3-HK concentration increased (Figure 4A). Quantification of the number of mitochondria present per cell showed no significant difference between untreated and treated samples; however, there was a significantly increased number of mitochondria present following treatment with 10 μM 3-HK when compared with 1 μM 3-HK (Figure 4B). There was also a trend towards an increased number of mitochondria present when samples were treated with 10 μM 3-HK relative to 0.1 μM 3-HK (P = 0.0698), but not when compared to the untreated control. In addition these data show a non-statistically significant but observable decrease in numbers of mitochondria present at 0.1 μM 3-HK compared to untreated cells.

**Figure 4:**
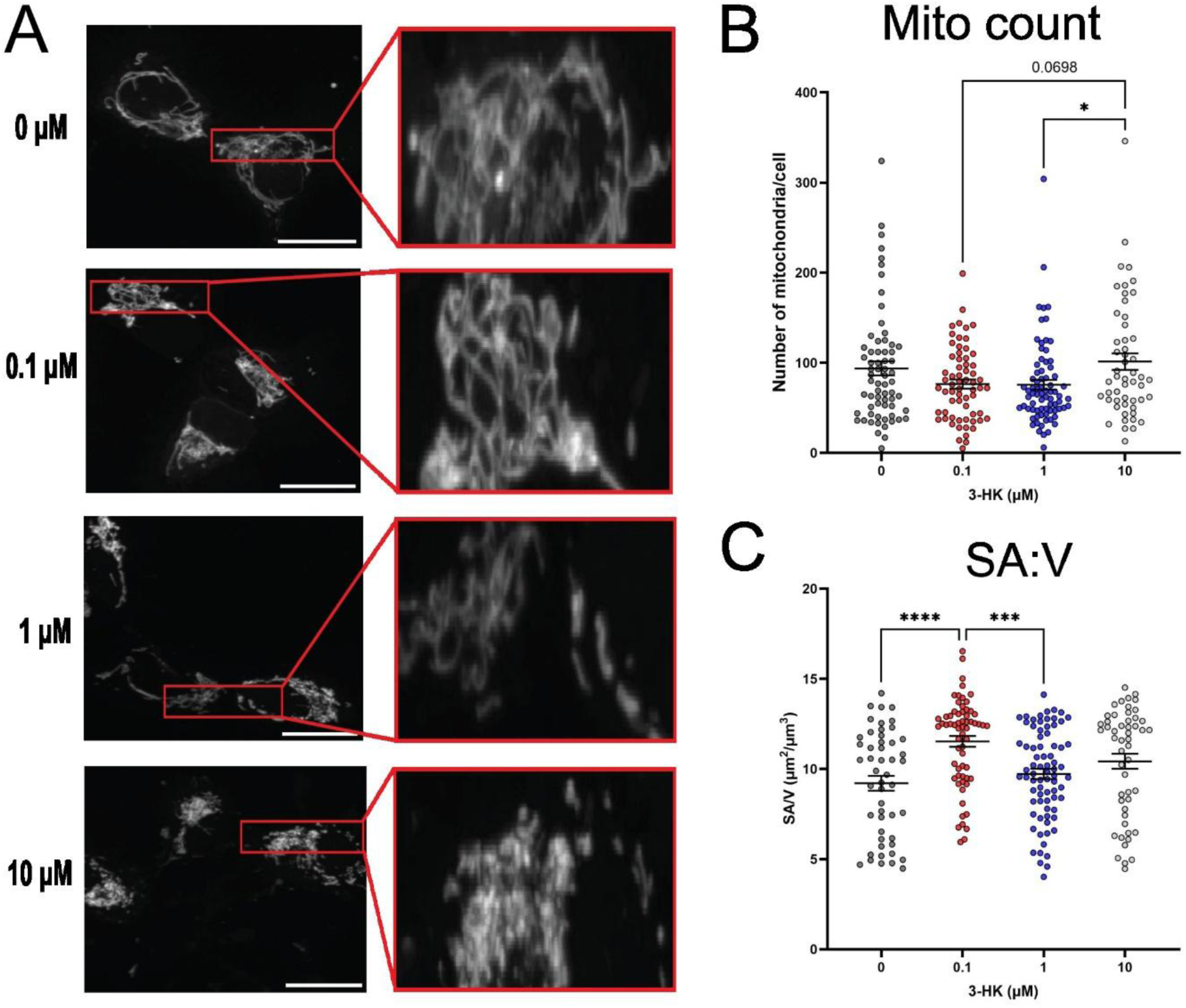
3-HK alters mitochondrial number and surface area to volume ratio (SA:V) in a biphasic manner. The mitochondrial networks of SH-SY5Y cells were stained using Mitoview^TM^ Green Dye and imaged by confocal microscopy following 24 hours exposure to 0, 0.1, 1 and 10 μM 3-HK. (A) Representative images per condition. Magnification X100. Scale bar = 20 μm. Red boxes indicate enlarged area. From analysis of 3D image stacks: (B) The number of mitochondria per cell. (C) SA:V (μm^2/^ μm^3^). Points represent a single cell and data are from 3 independent assays. One-way ANOVA with Tukey’s multiple comparisons test. Error bars indicate ± SEM. ****P < 0.0001. ***P < 0.001. *P < 0.05.

Following treatment with 0.1 μM 3-HK, there was a significant increase in the surface area to volume ratio (SA:V) of mitochondria present within the SH-SY5Y cells when compared with the drug-naïve samples (Figure 4C). There was no change in mitochondrial SA:V following treatment with ≥ 1 μM 3-HK, relative to the untreated control; however, there was a significant decrease in mitochondrial SA:V following treatment with 1 μM 3-HK when compared with that of 0.1 μM 3-HK. Again, the data suggest that the 0.1 μM 3-HK treated sample presents an unexpected mitochondrial phenotype that is not in line with treatments with higher concentrations of 3-HK. This phenomenon was observed in several other assessed metrics such as the total volume of mitochondria per cell (Figure 5A), the average volume per mitochondria (Figure 5B) and the average surface area per mitochondria (Figure 5C). However, no change in branch number or length was observed between any condition (Supp. Figure 3.)

**Figure 5:**
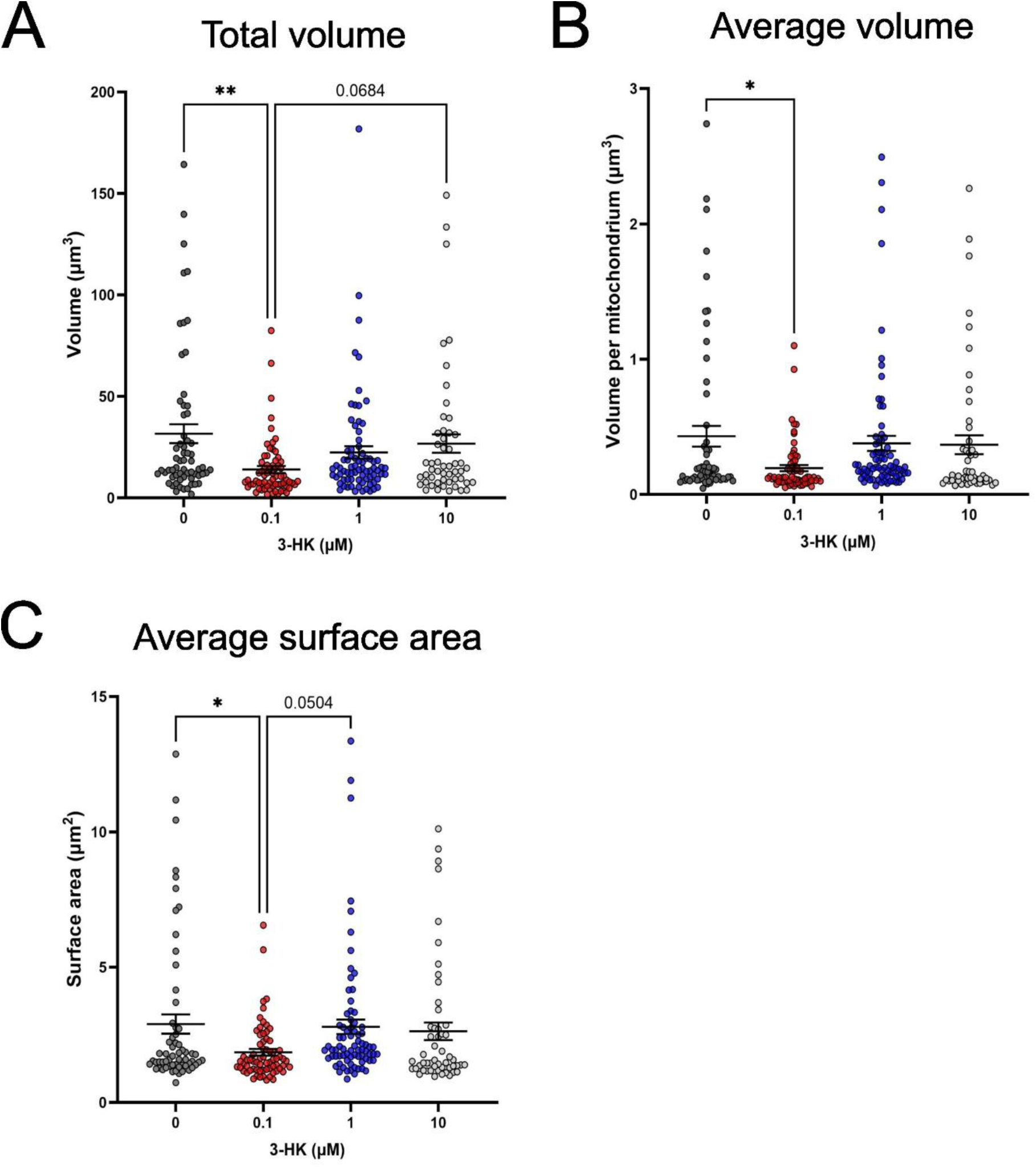
The mitochondrial networks of SH-SY5Y cells were stained using Mitoview^TM^ Green Dye and imaged by confocal microscopy following 24 hours exposure to 0, 0.1, 1 and 10 μM 3-HK. From analysis of 3D image stacks: (A) Total volume (μm^3^) of mitochondria per cell. (B) Average volume (μm^3^) per mitochondria. (C) Average surface area (μm^2^) per mitochondria. Points represent a single cell and data are from 3 independent assays. One-way ANOVA with Tukey’s multiple comparisons test. Error bars indicate ± SEM. **P < 0.01. *P < 0.05.

Mitochondrial dynamics rely heavily on cytoskeleton tracts which move mitochondria throughout the cell (Gatti *et al.,* 2025; Shi *et al.,* 2025). Therefore, filamentous actin (F-actin) density was assessed between conditions by fluorescence imaging; however, it was not found to differ (Supp. Figure 4).

### 3-HK treatment modulates the expression of mitochondrially relevant genes including *DNM1L*, *POLG*, *SIRT2* and *STRADA*

Finally, we explored transcriptional changes within SH-SY5Y cells following 3-HK treatment. We employed RNA-sequencing to identify differentially expressed genes (DEGs) linked to 3-HK treatment that might be specifically involved in the observed alterations in mitochondrial function and morphology. At all concentrations of 3-HK treatment (0.1 – 10 µM), a relatively modest number of genes (< 100) were found to be differentially expressed compared to both the untreated control and between all concentrations of 3-HK (Figure 6A, Tables S3 and S4). While only one DEG was common amongst all conditions, a small number of genes were shared between two conditions (19 between 0.1 vs 1 µM and 1 vs 10 µM; 12 between 0.1 vs 1 µM and 1 vs 10 µM; nine between 0.1 vs 10 µM and 1 vs 10 µM) (Supp. Figure 5; Supp. Table 5). Assessment of DEGs revealed that there was significantly decreased expression of the transcript NM_001278463, also known as *DNM1L,* observed at 0.1 µM 3-HK in comparison with the untreated condition (Figure 6B). *DNM1L* encodes for the protein Drp1, a key protein involved in mitochondrial fission. There was also significantly decreased expression of transcript NM_002693 (*POLG*) observed at 0.1 µM 3-HK in comparison with the untreated condition (Figure 6C). *POLG* encodes for the alpha subunit of DNA polymerase gamma, an enzyme essential for repairing and replicating mitochondrial DNA. Expression of NM_012237 (*SIRT2*) was significantly decreased at 0.1 µM 3-HK in comparison with the untreated condition but increased at 1 and 10 µM 3-HK in comparison with the 0.1 µM 3-HK condition (Figure 6D). *SIRT2* encodes for the NAD-dependent deacetylase sirtuin 2, a protein with links to many cellular processes, including mitochondrial homeostasis. Finally, there was a significantly decreased expression of NM_ 001003788 (*STRADA*) at 10 µM 3-HK in comparison with the 0.1 µM condition (Figure 6E). *STRADA* encodes for STE20-related kinase adaptor alpha, which is an upstream modulator of adenosine monophosphate-activated protein kinase (AMPK) which regulates mitochondrial biogenesis and function.

**Figure 6:**
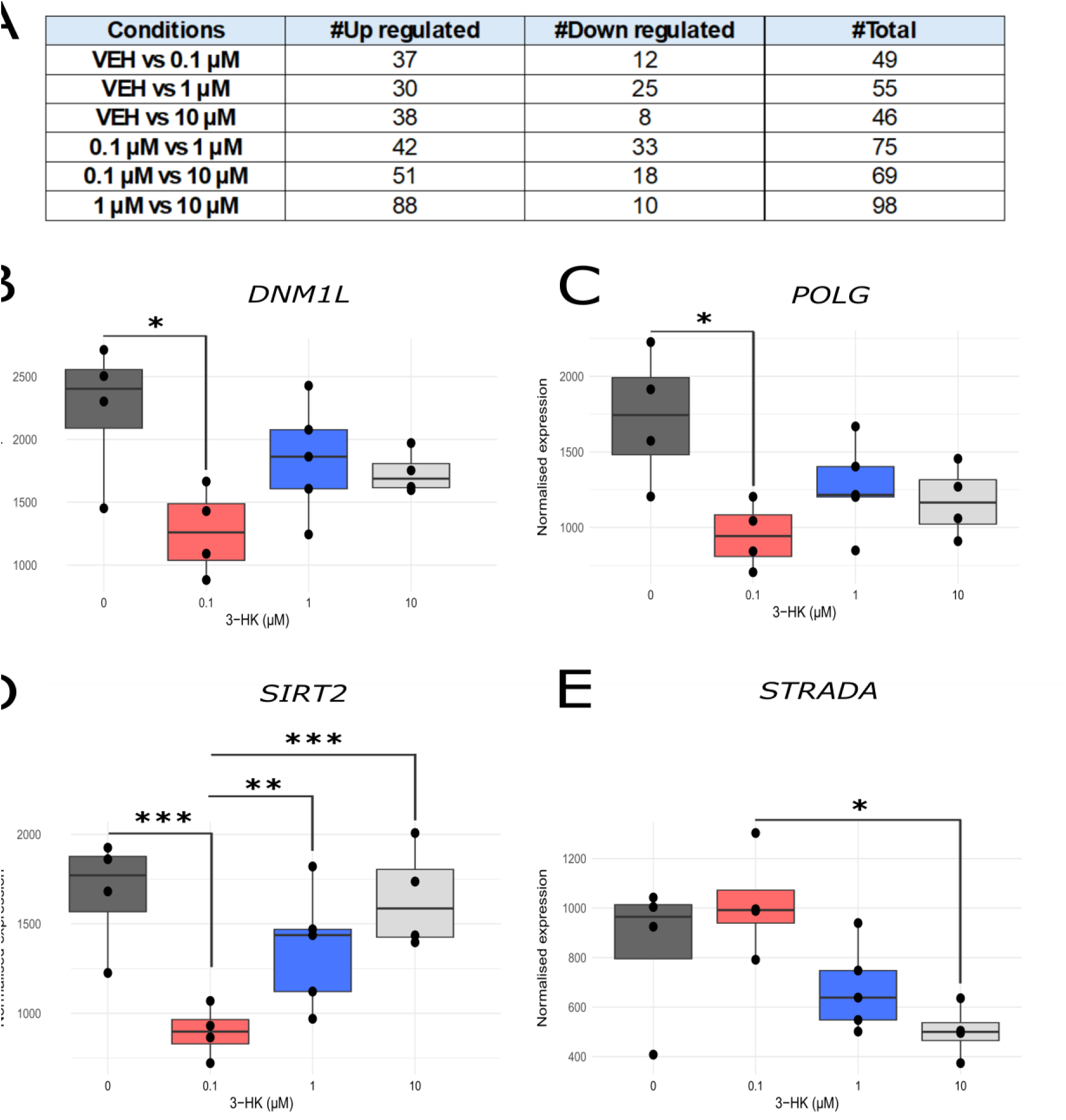
Transcriptomic profiling and differential gene expression analysis revealed altered expression of mitochondrially relevant genes following treatment with 3-HK. (A) DESeq2 results table showing the number of significantly up/down regulated differentially expressed transcripts in each experimental condition. *VEH = vehicle control.* (B) Normalised read counts of transcript NM_001278463 encoding for gene *DNM1L* when treated with 3-HK compared between conditions. (C) Normalised read counts of transcript NM_002693 encoding for gene *POLG* when treated with 3-HK compared between conditions. (D) Normalised read counts of transcript NM_012237 encoding for gene *SIRT2* when treated with 3-HK compared between conditions. (E) Normalised read counts of transcript NM_001003788 encoding for gene *STRADA* when treated with 3-HK compared between conditions. Each point represents one biological replicate. Interquartile range shown per condition. ***P < 0.001. **P < 0.01. *P < 0.05.

## Discussion

Neurons are the primary signalling cells in the brain but are critically influenced and supported by neighbouring non-neuronal cells, such as microglia, the brain’s resident immune cells (Frosch *et al.,* 2025; Marinelli *et al.,* 2019). Microglia play an import role in maintaining cerebral homeostasis as they constantly survey the microenvironment to regulate neurotransmitter signalling, clear debris and modulate inflammation (Bar & Barak, 2019; Herzog *et al.,* 2019; Li & Barres, 2018; Schafer *et al.,* 2013). In the event of cellular injury, stress or pathogen detection, microglial cells shift to an activated state, promoting neuroinflammation (Guo *et al.,* 2022). During this inflammatory response, neurotoxic factors are released by microglia which can cause neuronal dysfunction and ultimately lead to cellular death (Andreasen & Olsen, 1982; Olson & Miller, 2004). Notably, a key enzyme of the KP, KMO, is expressed in microglial cells, and activity of this enzyme is amplified during neuroinflammation (Connor *et al.,* 2008; Hirai *et al.,* 2010; Sathyasaikumar *et al.,* 2022). Expression of KMO is also increased in response to cytokines such as interleukin-1β, interferon-γ and tumour necrosis factor α (Esposito Soccoio *et al.,* 2026). This contributes to the elevated production of the KP metabolite 3-HK that has been shown to cause neuronal toxicity at supraphysiological levels in cell culture models (Buchanan *et al.,* 2023; Eastman & Guilarte, 1989; Okuda *et al.,* 1998; K. Wilson *et al.,* 2016). However, a dual role of 3-HK in the CNS has emerged where 3-HK has also been shown to have endogenous antioxidant properties mediated by its ability to donate electrons to contribute to redox reactions (Backhaus *et al.,* 2008; Christen *et al.,* 1990; Goshima *et al.,* 1986).

As inflammation stimulates microglial KMO activity, it is probable that 3-HK uptake by surrounding neurons is elevated in this instance. Due to the lack of clarity surrounding the role of 3-HK in neuronal physiology the aim of this study was to elucidate the cellular and molecular processes by which 3-HK elicits an effect with specific focus on the impact of increasing concentrations of this metabolite on neurons. This reflects the increased levels of 3-HK in the extracellular milieu during inflammation and in diseases of the CNS such as HD, PD and depressive disorder (Jasionowska *et al.,* 2024; Pearson & Reynolds, 1992; E. N. Wilson *et al.,* 2025). Additionally, we focused on mitochondrial function and morphology to examine how 3-HK, a known enhancer of H_2_O_2_ production (Okuda *et al.,* 1996), may modulate mitochondrial bioenergetics and dynamics. By characterising changes in mitochondrial function and structure under conditions that more closely reflect normal brain metabolism, this work aimed to clarify the role of 3-HK in modulating mitochondrial function and dynamics, and to provide insight into how alterations in the kynurenine pathway may contribute to early mitochondrial dysfunction in neurological disorders.

For our study, we employed 3-HK concentrations we determined to be physiologically relevant based on the literature. For example, a study by Pearson and Reynolds (1992) found that 3-HK was significantly elevated in the frontal cortex of post-mortem brain tissue from HD patients (from 0.15 ± 0.12 nmol/g to 0.41 ± 0.27 nmol/g). 3-HK was also significantly increased in the plasma of patients with depression compared with healthy individuals (from 8.92 ± 2.85 nM to 22.7 ± 3.48 nM) (Jasionowska *et al.,* 2024). Moreover, 3-HK levels were elevated in both the plasma and cerebral spinal fluid of PD patients when compared with a healthy control cohort (E. N. Wilson *et al.,* 2025).

Next, we assessed the activation of caspase-3/7 in neuroblastoma cells following treatment with 3-HK. Caspase-3/7 activation is a reliable indicator for the early stages of apoptosis and its significant increase in cells treated with 1 µM 3-HK suggests that despite the absence of change in cell viability measured at this concentration of 3-HK, at concentrations above healthy/normal 3-HK levels there is a drive in the system towards apoptosis. Both the extrinsic and intrinsic pathways of apoptosis promote caspase-3/7 activation (Lakhani *et al.,* 2006). The extrinsic pathway is triggered by external stimuli facilitating death receptor signalling, which 3-HK is not currently known to directly mediate (Green, 2022). However, the intrinsic apoptotic pathway is mitochondrially mediated, which 3-HK could directly initiate through production of H_2_O_2_ (Mustafa *et al.,* 2024; Okuda *et al.,* 1996, 1998). Therefore, it is feasible that caspase-3/7 activation observed with 3-HK reflects the ability of 3-HK to initiate early apoptotic events via increased mitochondrial H_2_O_2_ accumulation. Elevated ROS (as observed at 10 µM 3-HK) in the mitochondria would increase intracellular calcium, provoking cytochrome c release and alter expression of BCL-2 family proteins, ultimately activating caspase-3 and/or caspase-7 (Gutiérrez-Venegas *et al.,* 2015; Minzaghi *et al.,* 2023; Shoshan-Barmatz *et al.,* 2017).

We then examined the effect of 3-HK on cellular metabolic pathways by assessing redox capacity and ATP production. Cellular redox activity was found to be increased at 1 µM 3-HK, but not 10 µM 3-HK and only in comparison with 0.1 µM 3-HK and not the drug-naïve control, when measured by resazurin reduction to resorufin. Primarily this occurs in the mitochondria by complex I and II but resazurin can also be broken down by cytosolic enzymes such as oxidoreductases, dehydrogenases and redox active cofactors (Aleshin *et al.,* 2015; Zhang *et al.,* 2004). Interestingly, 3-HK had no effect on cellular ATP production. ATP is primarily derived from mitochondrial oxidative phosphorylation but can also be produced by glycolysis in the cytosol (Diaz-Ruiz *et al.,* 2011). Subsequently the effects of 3-HK directly on mitochondrial function were assessed using MitoSOX and revealed an increase in mitochondrial superoxide production at the highest tested 3-HK treatment (10 µM). Mitochondria superoxide is produced as a byproduct of the electron transport chain hence we sought to probe this further (Brand *et al.,* 2004). To do so, we employed the Seahorse Mito Stress assay where oxygen consumption rate of the cells following 3-HK treatment was investigated using inhibitors of the electron transport chain. However, respirometry measurements failed to show a difference between the untreated and 10 µM 3-HK conditions. Following assessment of mitochondrial network morphology by confocal imaging, and transcriptomic analyses, we believe that measurements may have been more informative had the lowest 3-HK treatment been assessed rather than the untreated condition.

Further probing the influence of 3-HK on mitochondrial functionality within our neuronal model revealed a biphasic effect on mitochondrial morphology. In our study, 0.1 µM 3-HK was the lowest concentration and was most relevant to normal homeostasis. The untreated condition is unrepresentative of human physiology as 3-HK is present at nanomolar concentrations as demonstrated in many studies (Aarsland *et al.,* 2015; Colín-González *et al.,* 2013; Curto *et al.,* 2016; Jasionowska *et al.,* 2024; E. N. Wilson *et al.,* 2025). Our data suggest that 3-HK at normal physiological levels may play a role in maintaining elongated mitochondrial networks, as there is a dramatic mitochondrial morphological difference observed in cells treated with the supposed persistent 3-HK levels found in healthy physiology. A significant decrease in mitochondrial total volume, average volume and average surface area was observed following treatment with 0.1 µM 3-HK, relative to the untreated control. Moreover, expression of *SIRT2,* which can be increased during neuroinflammation (Sola-Sevilla *et al.,* 2024) was significantly lower at 0.1 µM 3-HK, indicating a possible anti-inflammatory association at this concentration.

The mitochondrial SA:V ratio was significantly increased in cells treated with 0.1 µM 3-HK compared with the untreated control. This would typically imply a shift towards more, smaller mitochondria due to increased fission or decreased fusion (Bleck *et al.,* 2018; Botella *et al.,* 2023). However, we observed elongated networks with few individual mitochondria in cells treated with 0.1 µM 3-HK. Therefore, this increased SA:V combined with the non-fragmented nature of the mitochondria present in our samples suggest that the mitochondria present, whilst long, are much thinner than in the untreated condition. This is due to long, thin mitochondria in a constrained network having a higher SA:V as the volume decreases at a greater rate than the surface area (Preminger & Schuldiner, 2024; Rafelski, 2013; Schmiedl *et al.,* 1990). Mild stress can cause mitochondria to hyperfuse as a result of decreased fission or increased fusion (Gomes *et al.,* 2011; Rambold *et al.,* 2011; Tondera *et al.,* 2009; Yoon *et al.,* 2006). In our study, this may be due to the production of H_2_O_2_ by 3-HK causing low levels of cellular stress. This is in line with a study by Yoon et al,. (2006) who found that subcytotoxic doses of H_2_O_2_ promoted mitochondrial elongation in hepatocytes (Yoon *et al.,* 2006).

Transcriptomic analyses revealed that 3-HK altered the expression of multiple genes associated with mitochondrial health and function. The decreased expression of *DNM1L,* the gene which encodes for the mitochondrial fission regulator Drp1 (Lee *et al.,* 2004; Taguchi *et al.,* 2007), supports the hypothesis that mitochondria fission is dampened at normal physiological concentrations of 3-HK, in line with the elongated networks we observed at 0.1 µM during confocal imaging. The likelihood of decreased fission is further supported by the decreased *POLG* expression observed, indicating lower levels of mitochondrial DNA replication and repair (Graziewicz *et al.,* 2006). However, there was no statistically significant increase in *DNM1L* or *POLG* gene expression at higher 3-HK concentrations relative to either the untreated or lowest 3-HK condition. Therefore, it is unlikely that the increased mitochondrial fragmentation observed at higher concentrations of 3-HK was mediated through changes in Drp1 or POLG expression.

Our study provides evidence for a previously unknown role for 3-HK in regulating mitochondrial function and structure. This is likely through altered fission and fusion events due to modulated gene expression such as that of *DNML1* and our data suggests that subtle changes in KP metabolism regarding 3-HK may contribute to early mitochondrial dysfunction in neurological disease. Moreover, we highlight the importance of selecting the correct 3-HK concentration range for studies investigating metrics not overtly pertaining to cytotoxicity to mediate a more physiologically relevant understanding.

## Acknowledgements

We thank Nicolas Sylvius (Genomics Facility, University of Leicester) for assistance with RNA quality control and Kees Stratman (Advanced Imaging Facility, University of Leicester) for help with confocal microscopy. This work was supported by the Medical Research Council (MRC) and University of Leicester funded Integrated Midlands Partnership for Biomedical Training (IMPACT; grant number MR/N013913/1) and the National Institute of Mental Health (Silvio O. Conte, Center for Translational Mental Health Research – MH-103222).

