## Supplemental for "Physiological levels of 3-hydroxykynurenine alter mitochondrial function and morphology in neuronal cells"

### Supplementary data

**Supplementary Figure 1:** Characterisation of differentiated and undifferentiated SH-SY5Y cells. A) White light images of undifferentiated SH-SY5Y cells and cells differentiated for 11 days with 10 µM RA together with BDNF (12.5 ng/ml) for the final six days. Immunoblot analysis of (B) neuron specific enolase 2 and (C) MAP2 in undifferentiated and differentiated SH-SY5Y cells. (D) Representative images of undifferentiated SH-SY5Y cells following incubation with 3-HK (µM) for 24 hours. Mag = 20X. Scale bar = 100 µm.

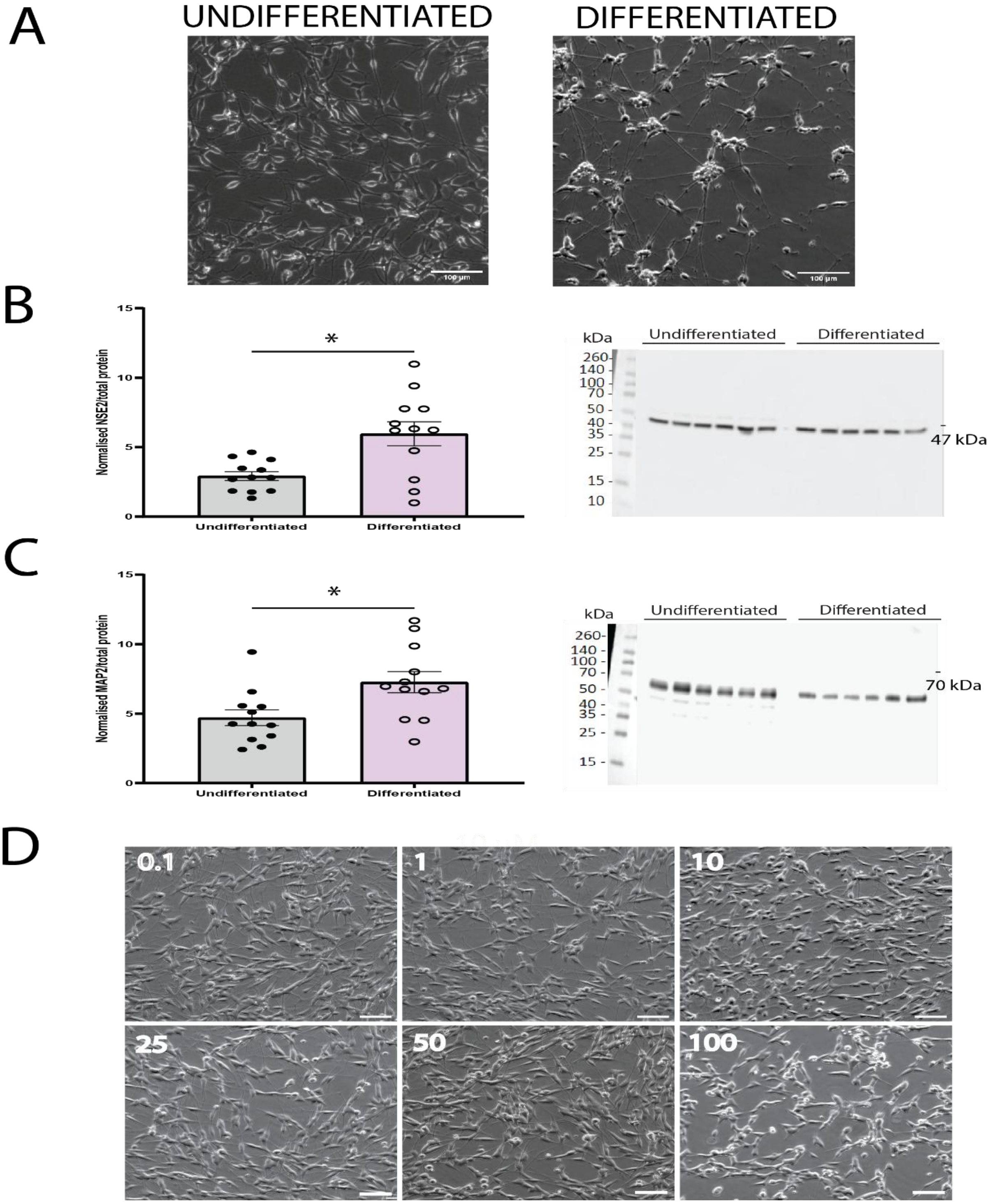

**Supplementary Table 1:** Kynurenine pathway and reference gene oligonucletides for qRT-PCR. Primers were purchased from Sigma-Aldrich. *3HAO = 3-hydroxyanthranilate oxidase. AADAT = Aminoadipate Aminotransferase. ACSMD = α-amino-β-carboxymuconate-ε-semialdehyde decarboxylase.* AFMID = arylformamidase. F = forward. IDO1 = *indoleamine 2,3-dioxygenase 1. IPO8 = importin 8. KMO = kynurenine 3-monooxygenase. CCBL1 = cysteine conjugate beta-lyase 1 (kynurenine aminotransferase 1). CCBL2 = cysteine conjugate beta-lyase 2 (kynurenine aminotransferase 3). KYNU = kynureninase. QPRT = Quinolinate phosphoribosyl transferase. qRT-PCR = quantitative reverse transcriptase polymerase chain reaction. R = reverse. Rplp0 = Ribosomal Protein Lateral Stalk Subunit P0. TDO2 = tryptophan 2,3-dioxygenase 2.*

| **Primer name** | **SEQUENCE (5'--> 3')** | **Description** |
|---|---|---|
| ***hIDO1-F*** | GCCAGCTTCGAGAAAGAGTTG | Forward primer for hIDO1 |
| ***hIDO1-R*** | ATCCCAGAACTAGACGTGCAA | Reverse primer for hIDO1 |
| ***hTDO2-F*** | AAGGTTGTTTCTCGGATGCAC | Forward primer for hTDO2 |
| ***hTDO2-R*** | TGTCATCGTCTCCAGAATGGAA | Reverse primer for hTDO2 |
| ***hAFMID-F*** | TGGGTTTCCCAAGCAAGGTTC | Forward primer for hAFMID |
| ***hAFMID-R*** | TCTGCTCCCAGTCGGACAA | Reverse primer for hAFMID |
| ***hCCBL1-F*** | CAGACTTTGCCGTGGAAGCCTT | Forward primer for hCCBL1 |
| ***hCCBL1-R*** | GCACATTCCTGAGCGGGTCTAT | Reverse primer for hCCBL1 |
| ***hAADAT-F*** | TGTCACATCTGGCAGCCAACAAG | Forward primer for hAADAT |
| ***hAADAT-R*** | GAATAAGCAGGTTCATCTAGGAGG | Reverse primer for hAADAT |
| ***hCCBL2-F*** | GTAGTGCTCCACTTACACGAGG | Forward primer for hCCBL2 |
| ***hCCBL2-R*** | GCTGCTTGAGTAACTGTCTCGAC | Reverse primer for hCCBL2 |
| ***hKMO-F*** | GAATGCGGGCTTTGAAGAC | Forward primer for hKMO |
| ***hKMO-R*** | ACAGGAAGACACAAACTAAGGT | Reverse primer for hKMO |
| ***hKYNU-F*** | GTTGGCTTTGATCTAGCACATGC | Forward primer for hKYNU |
| ***hKYNU-R*** | TGAAGGCACCAGCAATTCCTCC | Reverse primer for hKYNU |
| ***h3-HAO-F*** | CCTGAGACAGAATGTGGACGTG | Forward primer for h3-HAO |
| ***h3-HAO-R*** | CTTGTGTTCGCTCCCAGGCATA | Reverse primer for h3-HAO |
| ***hQPRT-F*** | GTGAAGGATAACCATGTGGTGGC | Forward primer for hQPRT |
| ***hQPRT-R*** | CTGCTGCATTCCACTTCCACCT | Reverse primer for hQPRT |
| ***hACMSD-F*** | GGTGCGAGAGAATTGCTGG | Forward primer for hACMSD |
| ***hACMSD-R*** | TGCTGGCAAGGTCGTTGTTT | Reverse primer for hACMSD |
| ***hIPO8-F*** | AGGATCAGAGGACAGCACTGCA | Forward primer for hIPO8 |
| ***hIPO8-R*** | AGGTGAAGCCTCCCTGTTGTTC | Reverse primer for hIPO8 |
| ***hRPLP0-F*** | GCTGCTGCCCGTGCTGGTG | Forward primer for hRPLP0 |
| ***hRPLP0-R*** | TGGTGCCCCTGGAGATTTTAGTGG | Reverse primer for hRPLP0 |

**Supplementary Table 2:** Kynurenine pathway enzymes constituently expressed in SH-SY5Y cells were detected using SYBR Green qRT-PCR. The assay was performed in triplicate. *NRT = reverse transcriptase null. AFMID = Arylformamidase. KYAT1 = Kynurenine aminotransferase 1. AADAT = Aminoadipate Aminotransferase. KYAT 3 = Kynurenine Aminotransferase 3. KMO = Kynurenine 3-Monooxygenase. IPO8 = importin 8. Rplp0 = Ribosomal Protein Lateral Stalk Subunit P0.*

| **Gene** | **Cell Phenotype** | **Ct Value** | |
|---|---|---|---|
| | | **Average** | **NRT** |
| **AFMID** | Non-differentiated | 27.0 | *Undetermined* |
| | Differentiated | 24.5 | *Undetermined* |
| **KYAT1** | Non-differentiated | 27.5 | *Undetermined* |
| | Differentiated | 27.2 | *Undetermined* |
| **AADAT** | Non-differentiated | 26.1 | *Undetermined* |
| | Differentiated | 27.0 | *Undetermined* |
| **KYAT3** | Non-differentiated | 22.0 | *Undetermined* |
| | Differentiated | 24.3 | *Undetermined* |
| **KMO** | Non-differentiated | 34.2 | 35.9 |
| | Differentiated | *Undetermined* | 31.1 |
| **IPO8** | Non-differentiated | 27.0 | *Undetermined* |
| | Differentiated | 23.9 | *Undetermined* |
| **RPLP0** | Non-differentiated | 19.5 | *Undetermined* |
| | Differentiated | 21.7 | *Undetermined* |

**Supplementary Figure 2:** Treatment of SH-SY5Y cells with 10 µM 3-HK had no effect on mitochondrial respiration when measured as oxygen consumption rate (OCR: pmol/minute µg/mL of protein) using the Seahorse bioanalyzer XFp. (A) Representative graph of the Seahorse Bioenergetic Mito Stress test (Agilent Biosciences) in cells treated with 10 µM 3-HK and control cells. (B) Basal respiration, (C) ATP-linked respiration, (D) non-mitochondrial respiration and (E) coupling efficiency (%) (ATP-linked respiration rate/ basal respiration rate ×100) were quantified. Points represent the average value per replicate from 3 independent assays. Unpaired Student's t-test. Error bars indicate ± SEM.

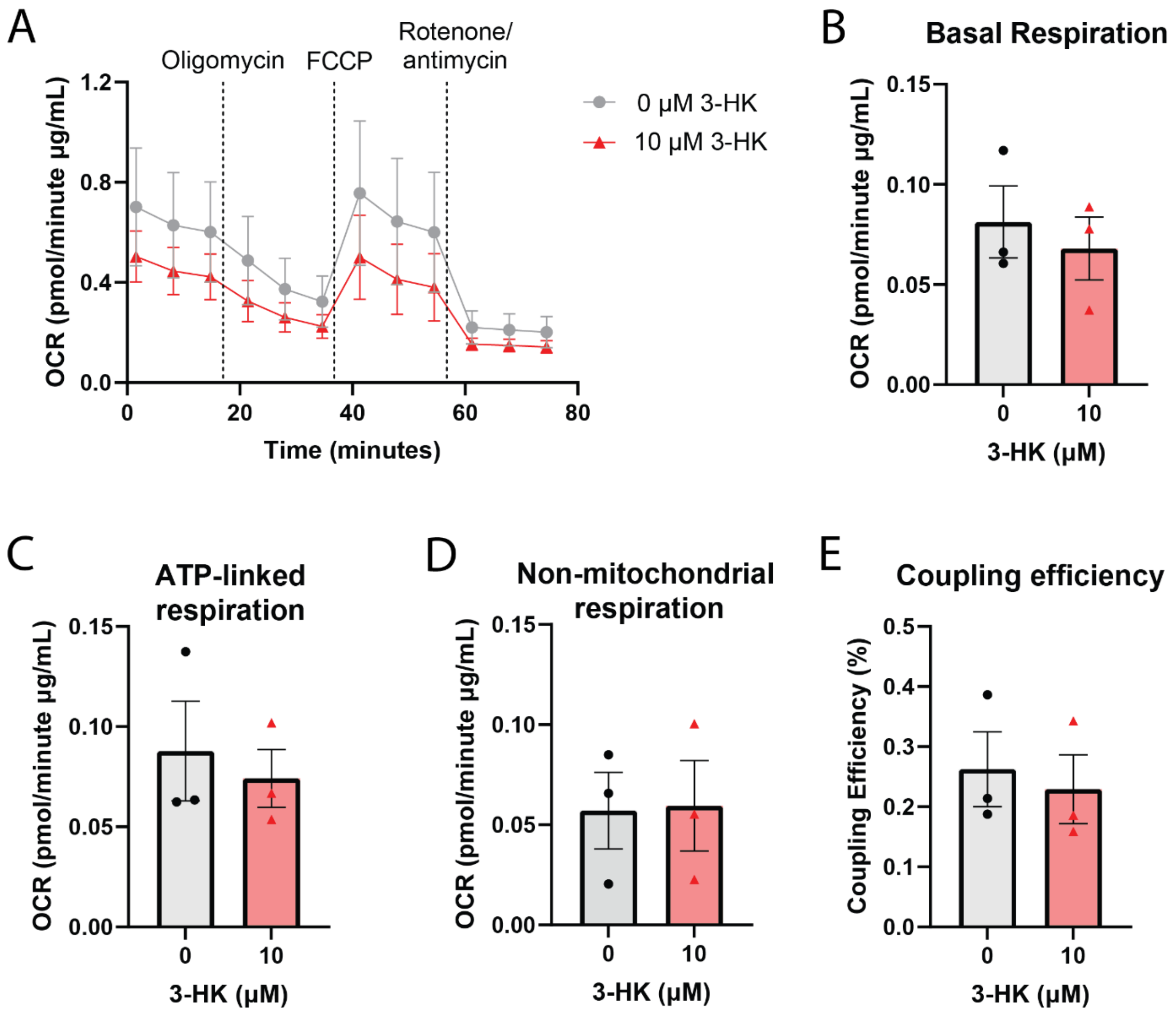

**Supplementary Figure 3:** The mitochondrial networks of SH-SY5Y cells were stained using Mitoview™ Green Dye and imaged by confocal microscopy for analysis following 24 hours exposure to 0, 0.1, 1 and 10 µM 3-HK. From maximum projection images: (A) The number of branches per mitochondria. (B) The average mitochondrial branch length (µm) per cell. Points represent a single cell and data is from 3 independent assays. Error bars indicate ± SEM.

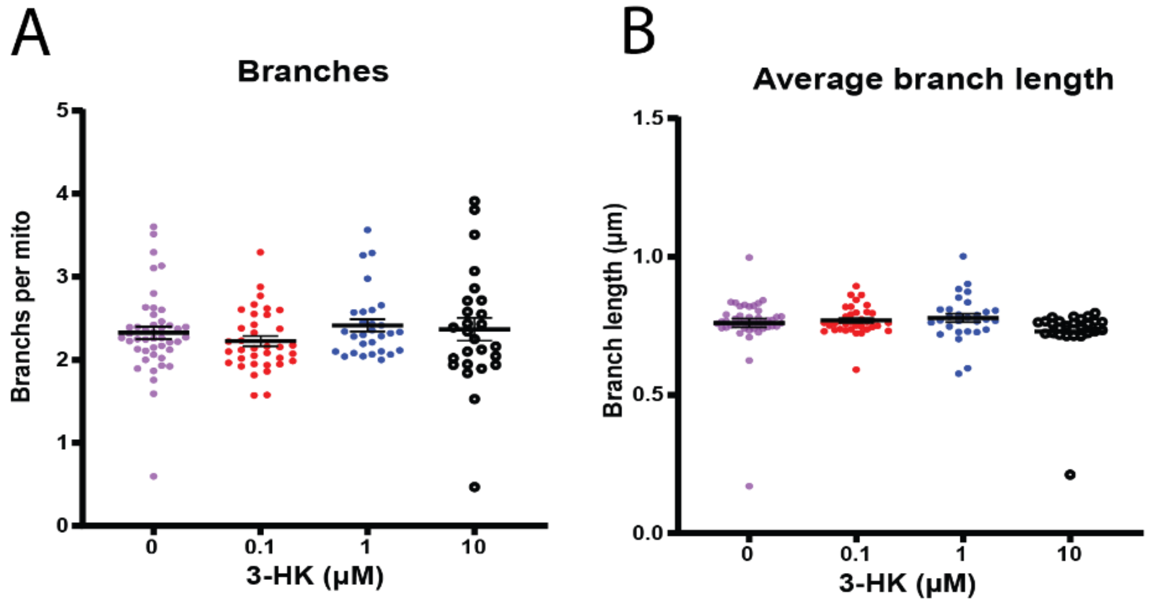

**Supplementary Figure 4:** F-actin staining of SH-SY5Y cells was unchanged following exogenous 3-HK treatment. Fluorescence intensity (AU) of Phalloidin-594 staining was measured via confocal microscopy to quantify F-actin. (A) No change in F-actin staining was observed at any tested concentration of 3-HK (µM) in comparison with the vehicle control. Points each represent a single cell and data is from 3 independent assays. One-way ANOVA. Error bars indicate ± SEM. (B) Representative images per condition. Magnification x100. Scale bar = 20 µm. *VEH = vehicle control.*

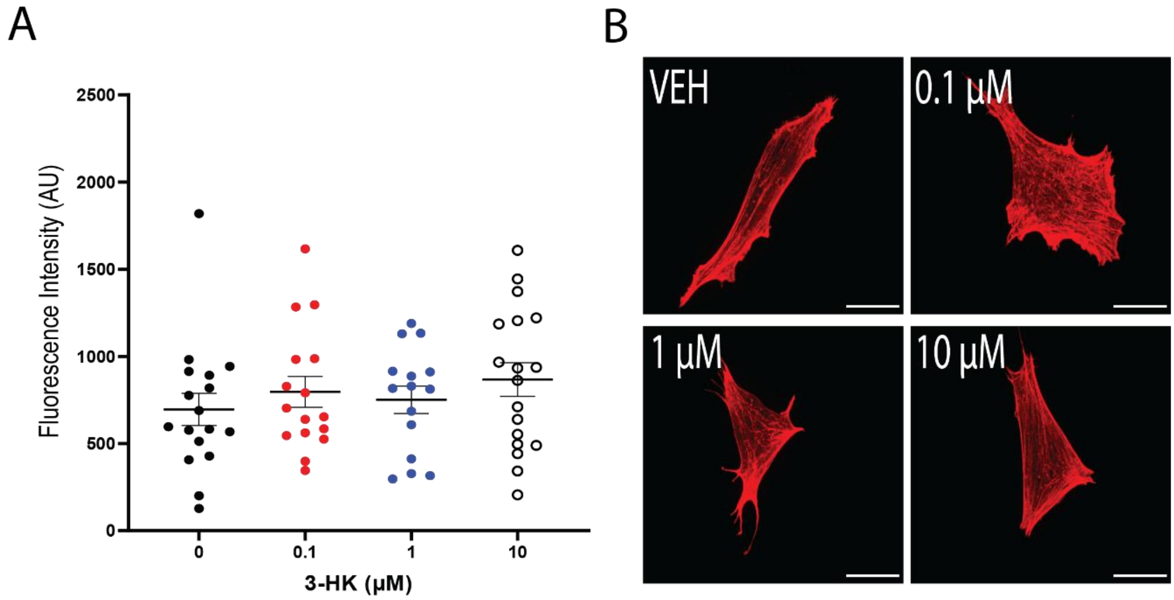

**Supplementary Table 3:** DESeq2 results table showing significantly up regulated differentially expressed transcripts (DETs) in each condition. Each transcript is presented by both its unique RefSeq transcript ID and the related common gene ID/symbol, where possible. Transcripts not mapped to a specific gene ID are shown as 'None'. The log2FoldChange (L2FC) and adjusted P-value (padj) are noted. Transcripts are ordered by descending log2FoldChange.

| UP REGULATED | | | |
|---|---|---|---|
| **TRANSCRIPT ID** | **GENE SYMBOL** | **L2FC** | **PADJ** |
| **VEHICLE CONTROL VS 0.1 µM 3-HK** | | | |
| NM_001256270 | KIF22 | 9.738339 | 0.001832 |
| NM_024767 | DLC1 | 7.997934 | 0.002549 |
| NM_001278393 | USP44 | 5.554421 | 2.28E-06 |
| NM_001040110 | NRF1 | 4.986878 | 0.005566 |
| NM_001284338 | NEDD4 | 4.256312 | 0.015145 |
| NM_022963 | FGFR4 | 3.956237 | 0.014643 |
| NM_001204459 | TNFRSF19 | 3.744702 | 0.030273 |
| NM_001256163 | BIRC2 | 3.358783 | 1.41E-06 |
| NR_027489 | MTHFSD | 3.28713 | 0.006826 |
| NM_001282585 | MXRA8 | 2.672457 | 0.013677 |
| NM_001166686 | PFKM | 2.629228 | 0.011187 |
| NM_001161533 | None | 2.441415 | 0.00011 |
| NM_001163436 | TBCK | 2.352795 | 0.003614 |
| NR_105052 | PSD2-AS1 | 2.038559 | 0.035616 |
| NM_015493 | KANK2 | 1.992757 | 0.002107 |
| NM_001191324 | RNF138 | 1.859071 | 0.042656 |
| NM_001254749 | RGS5 | 1.691107 | 2.45E-05 |
| NM_001009812 | LBX2 | 1.664739 | 0.038129 |
| NM_001256879 | POLD2 | 1.593209 | 0.03099 |
| NM_032853 | PWWP3A | 1.513908 | 1.71E-07 |
| NM_001042595 | TMEM91 | 1.429053 | 0.018702 |
| NM_134426 | SLC26A6 | 1.368685 | 0.036484 |
| NM_001097594 | XAGE1A | 1.329517 | 0.015363 |
| NR_027167 | SNHG29 | 1.289702 | 0.004533 |
| NM_001040454 | SLC26A6 | 1.244248 | 0.006826 |
| NM_004772 | NREP | 1.201608 | 8.92E-06 |
| NM_001286672 | AASDH | 1.110469 | 0.002512 |
| NR_130944 | WDR73 | 1.081557 | 0.02321 |
| NM_001080427 | None | 1.008411 | 0.007821 |
| NM_003250 | THRA | 0.930419 | 0.007246 |
| NR_027504 | MST1P2 | 0.814836 | 0.009405 |
| NM_006531 | IFT88 | 0.810251 | 0.002496 |
| NR_002190 | SUMO1P3 | 0.798795 | 0.017907 |
| NR_024166 | ZNF205-AS1 | 0.792977 | 0.04904 |
| NR_104307 | ULK3 | 0.773158 | 0.02042 |
| NM_002982 | CCL2 | 0.686678 | 0.020731 |
| NM_001098577 | RPL31 | 0.653554 | 0.008703 |
| NR_038879 | PAXBP1-AS1 | 0.632831 | 0.013722 |
| NM_003260 | TLE2 | 0.572646 | 0.004957 |
| **VEHICLE CONTROL VS 1 µM 3-HK** | | | |
| NM_033668 | ITGB1 | 6.920805 | 0.000833 |
| NM_001284274 | HECTD2 | 5.748252 | 0.019864 |
| NM_001164824 | SMIM12 | 5.538854 | 0.001602 |
| NM_000704 | ATP4A | 5.465517 | 0.002114 |
| NR_040712 | None | 5.280932 | 0.015731 |

| | | | |
|---|---|---|---|
| NR_024125 | ATP1A1-AS1 | 4.796088 | 0.028071 |
| NM_001261438 | ETV4 | 4.623017 | 0.004446 |
| NM_001297778 | NMNAT1 | 4.473952 | 0.000337 |
| NM_001282660 | MRTFA | 4.391425 | 4.61E-16 |
| NM_001303108 | PGAP4 | 4.125634 | 0.01968 |
| NM_001287044 | VEGFA | 3.995145 | 0.021994 |
| NM_001079877 | RASA4 | 3.901976 | 4.36E-05 |
| NM_174963 | ST3GAL3 | 3.764657 | 0.000381 |
| NM_001166686 | PFKM | 3.011991 | 0.01281 |
| NR_002715 | RN7SL1 | 2.979518 | 0.003567 |
| NM_001282568 | ZCCHC17 | 2.751469 | 0.013743 |
| NM_001242701 | CAMTA1 | 2.695539 | 0.005622 |
| NM_001009812 | LBX2 | 2.181117 | 0.011696 |
| NR_027260 | RN7SL2 | 2.174626 | 0.039876 |
| NM_001454 | FOXJ1 | 2.026516 | 0.004243 |
| NM_174971 | ST3GAL3 | 2.011957 | 0.030466 |
| NM_022805 | SNRPN | 1.592899 | 0.029847 |
| NM_199191 | BABAM2 | 1.440381 | 0.001472 |
| NM_003099 | SNX1 | 1.325315 | 0.021607 |
| NM_001291018 | PDE6D | 1.308704 | 0.043573 |
| NR_029390 | None | 1.248058 | 0.016125 |
| NR_110373 | LINC01159 | 1.167409 | 0.02754 |
| NM_001012973 | PLAC9 | 0.895829 | 0.045922 |
| NM_003260 | TLE2 | 0.730584 | 0.002063 |
| NR_036487 | FGF14-AS2 | 0.563224 | 0.015484 |
| **VEHICLE CONTROL VS 10 μM 3-HK** | | | |
| NM_001102664 | EPN2 | 9.92656 | 0.001089 |
| NM_015350 | LRRC8B | 5.90399 | 0.00944 |
| NM_017787 | WBP1L | 5.698395 | 0.036647 |
| NM_001282886 | EIF4E3 | 5.574405 | 0.000974 |
| NM_001278393 | USP44 | 5.528973 | 3.01E-06 |
| NM_001199014 | STPG1 | 4.892734 | 0.000981 |
| NR_003286_2 | None | 4.779938 | 0.001333 |
| NM_001100603 | KDELR2 | 4.512632 | 0.038057 |
| NM_001271960 | SLC39A1 | 4.460067 | 0.001714 |
| NM_001290307 | CTNNA1 | 4.046996 | 0.037501 |
| NM_001174097 | LDHB | 3.696275 | 0.031055 |
| NM_001256335 | PTGES2 | 3.595795 | 2.73E-05 |
| NM_015520 | MAGI1 | 3.269136 | 0.01912 |
| NM_001257361 | AMPD2 | 3.112858 | 0.003104 |
| NM_001282568 | ZCCHC17 | 2.807842 | 0.002353 |
| NM_001286109 | CLN3 | 2.4998 | 0.016657 |
| NM_001307951 | LPXN | 2.33095 | 0.046679 |
| NM_032853 | PWWP3A | 2.274606 | 7.40E-16 |
| NM_001009812 | LBX2 | 2.245881 | 0.002176 |
| NM_003169 | SUPT5H | 2.076136 | 0.003114 |
| NM_006317 | BASP1 | 1.858747 | 0.000635 |
| NM_001080427 | None | 1.81309 | 4.12E-08 |
| NM_001243204 | ECSIT | 1.715635 | 0.040649 |
| NM_001161533 | None | 1.650452 | 0.023048 |
| NR_046096 | CPEB1-AS1 | 1.62884 | 0.017078 |
| NR_110542 | PIK3IP1-DT | 1.598313 | 0.007181 |
| NM_172095 | CATSPER2 | 1.40655 | 1.58E-05 |
| NM_001243738 | RGL2 | 1.353189 | 0.004706 |

| | | | |
|---|---|---|---|
| NM_001282583 | MXRA8 | 1.296908 | 0.013626 |
| NR_003574 | ABCA17P | 1.251936 | 0.007086 |
| NM_020877 | DNAH2 | 1.217342 | 0.00842 |
| NM_001282497 | ZNF343 | 1.097727 | 0.0313 |
| NR_110220 | LINC01237 | 0.979032 | 0.048142 |
| NM_022468 | MMP25 | 0.883683 | 0.037778 |
| NM_014741 | ATG13 | 0.761813 | 0.04611 |
| NM_015833 | ADARB1 | 0.753944 | 0.022722 |
| NM_001300927 | PGPEP1 | 0.62556 | 0.048542 |
| NM_001282494_1 | None | 0.59094 | 0.003898 |
| **0.1 μM 3-HK VS 1 μM 3-HK** | | | |
| NM_001277335 | RASA4B | 14.66796 | 8.81E-12 |
| NM_052985 | IFT122 | 9.89781 | 3.38E-05 |
| NM_001204887 | RAB43 | 9.408757 | 0.001838 |
| NM_001303108 | PGAP4 | 7.189597 | 2.52E-05 |
| NM_001284274 | HECTD2 | 6.6406 | 0.00777 |
| NM_033668 | ITGB1 | 6.479809 | 0.001687 |
| NM_001164824 | SMIM12 | 6.022465 | 0.000528 |
| NR_040712 | CHRAC1 | 5.12465 | 0.01854 |
| NM_001297778 | NMNAT1 | 4.494715 | 0.0003 |
| NM_001282660 | MRTFA | 4.387944 | 4.92E-16 |
| NM_001206929 | AGER | 4.188927 | 0.03536 |
| NM_001261438 | ETV4 | 3.718227 | 0.02305 |
| NM_174963 | ST3GAL3 | 3.652346 | 0.000546 |
| NM_001278205 | None | 3.589867 | 0.024454 |
| NM_201595 | GTF2A1 | 3.574857 | 0.025728 |
| NM_001142310 | TMEM169 | 3.153783 | 0.029249 |
| NM_001170580 | HHAT | 3.07133 | 0.027967 |
| NM_174971 | ST3GAL3 | 2.963488 | 0.00093 |
| NM_001173482 | CRBN | 2.942856 | 0.03165 |
| NR_001445 | RN7SK | 2.82016 | 0.030828 |
| NM_001273 | CHD4 | 2.785769 | 5.42E-08 |
| NR_002715 | RN7SL1 | 2.591334 | 0.010741 |
| NR_002569 | SCARNA9 | 2.471803 | 0.003653 |
| NM_001013406 | KRIT1 | 2.369748 | 0.014035 |
| NM_020734 | RIMKLB | 2.26333 | 0.02322 |
| NM_001105540 | DGKZ | 1.9011 | 0.000196 |
| NM_003455 | ZNF202 | 1.884836 | 1.44E-05 |
| NM_001813 | CENPE | 1.801537 | 0.033902 |
| NM_001205179 | ALKBH2 | 1.781041 | 0.014898 |
| NM_001168222 | TBC1D17 | 1.705659 | 0.009661 |
| NM_001282652 | STIP1 | 1.645486 | 0.006908 |
| NM_001278189 | TPM3 | 1.588036 | 7.01E-07 |
| NM_052870 | SNX18 | 1.575437 | 0.00898 |
| NM_001454 | FOXJ1 | 1.525945 | 0.029821 |
| NM_003099 | SNX1 | 1.410926 | 0.012915 |
| NR_033489 | ABHD16A | 1.202027 | 0.026276 |
| NR_051960 | FALEC | 1.090007 | 0.039852 |
| NM_001286582 | PHRF1 | 1.079649 | 0.047857 |
| NM_001024809 | RARA | 1.019037 | 0.010329 |
| NM_001267783 | AMBRA1 | 0.909821 | 0.001532 |
| NR_073599 | None | 0.893083 | 0.005928 |
| NM_181727 | SPATA12 | 0.840312 | 0.038043 |
| NM_012237 | SIRT2 | 0.712758 | 0.00898 |

| | | | |
|---|---|---|---|
| NM_024033 | CYREN | 0.682922 | 0.023726 |
| **0.1 μM 3-HK VS 10 μM 3-HK** | | | |
| NM_001277335 | RASA4B | 14.6841 | 3.75E-15 |
| NM_017787 | WBP1L | 11.43479 | 0.000126 |
| NM_001098493 | ZNF419 | 10.10302 | 2.22E-07 |
| NM_001098496 | ZNF419 | 9.618508 | 1.68E-06 |
| NR_103455 | ZSCAN16-AS1 | 8.910362 | 3.00E-07 |
| NM_001277947 | ZNF83 | 7.941939 | 0.00268 |
| NM_178173 | IHO1 | 6.298739 | 0.01756 |
| NR_003286_2 | None | 5.049326 | 0.000478 |
| NM_001172225 | ZNF540 | 4.869123 | 0.018021 |
| NM_001303108 | PGAP4 | 4.846815 | 0.000821 |
| NM_001278926 | PPWD1 | 4.553677 | 0.033 |
| NM_202758 | ENOSF1 | 4.5324 | 0.016696 |
| NM_001193369 | DIDO1 | 3.920519 | 0.0001 |
| NR_040053 | RNF41 | 3.759373 | 0.000511 |
| NM_001197079 | IFRD1 | 3.694166 | 0.02182 |
| NM_001282719 | LDAH | 3.619675 | 0.004744 |
| NM_001267818 | OSTC | 3.607314 | 0.000688 |
| NR_033836 | PIGA | 3.602479 | 0.008018 |
| NM_001278205 | None | 3.219562 | 0.018096 |
| NM_001173482 | CRBN | 3.20516 | 0.004686 |
| NR_037649 | CCNT2 | 3.181526 | 0.007023 |
| NM_015520 | MAGI1 | 3.131458 | 0.02244 |
| NM_001145548 | ZDHHC7 | 3.089417 | 0.020209 |
| NM_001271960 | SLC39A1 | 2.986281 | 0.044194 |
| NM_153047 | FYN | 2.802812 | 0.010196 |
| NM_181527 | NAA20 | 2.557755 | 0.01316 |
| NM_001813 | CENPE | 2.493094 | 0.000205 |
| NM_001145287 | OPRM1 | 2.456373 | 0.007175 |
| NM_001273 | CHD4 | 2.413364 | 5.37E-09 |
| NM_001256335 | PTGES2 | 2.105594 | 0.025784 |
| NM_001160390 | TRPT1 | 1.996691 | 0.006643 |
| NM_001205179 | ALKBH2 | 1.98907 | 0.000992 |
| NM_001105540 | DGKZ | 1.9581 | 1.38E-06 |
| NM_006317 | BASP1 | 1.848278 | 0.000588 |
| NM_001278189 | TPM3 | 1.656047 | 8.01E-11 |
| NM_003169 | SUPT5H | 1.63318 | 0.026997 |
| NM_001243738 | RGL2 | 1.539372 | 0.000694 |
| NR_034121 | CKMT2-AS1 | 1.403649 | 0.042249 |
| NM_001145011 | C16orf96 | 1.17382 | 0.044602 |
| NM_001204368 | MGST2 | 1.167481 | 0.038044 |
| NR_120577 | ESAM-AS1 | 1.091333 | 0.000932 |
| NR_110220 | LINC01237 | 1.012859 | 0.034283 |
| NM_001282723 | LDAH | 1.003243 | 0.017433 |
| NR_026974 | ZNF252P-AS1 | 0.95234 | 0.030937 |
| NM_199334 | THRA | 0.946991 | 0.001362 |
| NM_002285 | AFF3 | 0.892093 | 0.030675 |
| NM_198395 | G3BP1 | 0.843925 | 0.007297 |
| NM_001285450 | PDXDC1 | 0.813798 | 0.030937 |
| NM_001301072 | MAP3K4 | 0.802075 | 0.038854 |
| NM_012237 | SIRT2 | 0.783508 | 0.000416 |
| NM_002941 | ROBO1 | 0.730228 | 0.000716 |
| NM_001204747 | RFC1 | 0.721332 | 0.014222 |

| | | | |
|---|---|---|---|
| NR_002473 | None | 0.678943 | 0.023574 |
| NR_040013 | LOC644554 | 0.610662 | 0.013787 |
| **1 μM 3-HK VS 10 μM 3-HK** | | | |
| NM_001135651 | EIF2AK2 | 14.31435 | 9.61E-11 |
| NM_024772 | ZMYM1 | 13.10537 | 1.32E-14 |
| NM_001194955 | MATR3 | 11.39406 | 0.000107 |
| NM_001199014 | STPG1 | 10.5672 | 2.06E-05 |
| NR_040762 | ZIC4 | 9.613276 | 2.58E-07 |
| NM_001310339 | MGME1 | 9.535294 | 5.87E-07 |
| NR_033836 | PIGA | 9.16116 | 8.27E-05 |
| NM_194454 | KRIT1 | 8.439637 | 1.55E-14 |
| NM_006661 | PDE10A | 8.185792 | 3.19E-07 |
| NM_001098496 | ZNF419 | 8.142011 | 0.000895 |
| NM_001172225 | ZNF540 | 7.799394 | 0.006933 |
| NM_005520 | HNRNPH1 | 6.839359 | 7.02E-07 |
| NM_001288632 | FAM222B | 6.833451 | 0.00123 |
| NM_014432 | IL20RA | 6.562071 | 6.23E-08 |
| NM_024515 | WDR25 | 5.88177 | 4.41E-05 |
| NM_175735 | LYG2 | 5.632041 | 0.002641 |
| NR_110156 | LINC01798 | 5.495469 | 0.005721 |
| NR_003286_2 | None | 5.481979 | 0.001656 |
| NM_148955 | SNX1 | 5.393912 | 5.02E-08 |
| NM_001098493 | ZNF419 | 5.33323 | 0.004464 |
| NM_001206799 | PKM | 5.117624 | 2.00E-05 |
| NM_001164747 | RASSF8 | 4.999121 | 0.00071 |
| NM_001204171 | MDM4 | 4.682813 | 1.54E-10 |
| NM_017996 | DET1 | 4.24968 | 6.56E-07 |
| NM_005734 | HIPK3 | 4.191239 | 2.71E-05 |
| NM_181482 | LDLRAD4 | 4.081661 | 0.000714 |
| NR_040585 | STAG3L4 | 4.048867 | 0.001441 |
| NM_001256335 | PTGES2 | 3.694882 | 0.000656 |
| NM_004361 | CDH7 | 3.626062 | 0.000432 |
| NM_001286837 | RNASEH1 | 3.02481 | 0.000825 |
| NM_001278174 | ZNF33A | 2.955388 | 0.020557 |
| NM_001290259 | PHF3 | 2.903828 | 0.003864 |
| NM_001178011 | CDC45 | 2.865359 | 0.002805 |
| NM_001085377 | MCC | 2.778641 | 0.023358 |
| NM_001142327 | DMTF1 | 2.778022 | 0.002109 |
| NM_001005741 | GBA | 2.593427 | 1.82E-05 |
| NR_125792 | LINC01291 | 2.500448 | 0.038646 |
| NM_001114617 | MGAT1 | 2.391266 | 5.30E-06 |
| NM_001243374 | CLCN3 | 2.365504 | 0.009235 |
| NM_203401 | STMN1 | 2.270622 | 0.028262 |
| NM_197966 | BID | 2.227611 | 2.80E-05 |
| NM_001130849 | CAB39 | 2.211567 | 0.042984 |
| NM_001134774 | KLC2 | 2.198945 | 0.001911 |
| NM_001286589 | AIG1 | 2.178339 | 0.007888 |
| NM_024731 | KLHL36 | 2.048586 | 0.003977 |
| NR_024514 | ADAMTS13 | 2.00693 | 0.049828 |
| NM_001170780 | PABIR3 | 1.936016 | 0.03442 |
| NM_006317 | BASP1 | 1.857437 | 0.005074 |
| NM_032853 | PWWP3A | 1.787955 | 5.87E-07 |
| NM_203505 | G3BP2 | 1.775836 | 0.005852 |
| NM_000093 | COL5A1 | 1.739793 | 4.77E-05 |

| | | | |
|---|---|---|---|
| NM_022372 | MLST8 | 1.685967 | 0.011742 |
| NM_001301819 | ZNF202 | 1.676746 | 6.60E-06 |
| NM_003671 | CDC14B | 1.674478 | 0.008302 |
| NM_003169 | SUPT5H | 1.670962 | 0.037517 |
| NM_012472 | DNAAF11 | 1.598646 | 0.000522 |
| NM_001014 | RPS10 | 1.570797 | 0.008306 |
| NM_001455 | FOXO3 | 1.511315 | 0.025897 |
| NM_031501 | PCDHA5 | 1.316989 | 0.012225 |
| NM_001199746 | HOXD8 | 1.284557 | 0.008117 |
| NM_000333 | ATXN7 | 1.253063 | 0.042479 |
| NM_138559 | BCL11A | 1.219127 | 0.02651 |
| NR_125729 | LINC00680 | 1.196582 | 0.014243 |
| NR_110099 | SNHG21 | 1.190132 | 0.010027 |
| NM_024867 | SPEF2 | 1.179244 | 0.009566 |
| NM_022107 | GPSM3 | 1.178587 | 0.002272 |
| NR_073395 | FAM90A25P | 1.172752 | 0.014941 |
| NM_018850 | ABCB4 | 1.162026 | 0.002582 |
| NM_001076785 | SLC7A6 | 1.120457 | 0.021053 |
| NM_024877 | CCNP | 1.109266 | 0.047639 |
| NM_001080427 | None | 1.094523 | 0.008484 |
| NM_180699 | SNRNP35 | 1.081142 | 0.021874 |
| NM_001285450 | PDXDC1 | 1.070917 | 0.01132 |
| NM_001267574 | EIF3C | 1.048598 | 0.005966 |
| NM_021038 | MBNL1 | 1.035235 | 0.032834 |
| NR_122109 | CYP51A1-AS1 | 1.034508 | 0.030416 |
| NM_001127361 | RNF19B | 1.009333 | 0.041696 |
| NM_001172677 | ZNF607 | 0.990643 | 0.007678 |
| NM_001199355 | RPL17-C18orf32 | 0.979912 | 0.008693 |
| NM_005888 | SLC25A3 | 0.969299 | 0.013333 |
| NM_001278217 | CDK5RAP3 | 0.94933 | 0.04678 |
| NM_022757 | CCDC14 | 0.923131 | 0.043744 |
| NM_020831 | MRTFA | 0.90457 | 1.40E-07 |
| NR_040013 | LOC644554 | 0.871154 | 0.003091 |
| NM_007222 | ZHX1 | 0.819201 | 0.009146 |
| NM_001668 | ARNT | 0.739339 | 0.036353 |
| NM_153255 | MCM9 | 0.685808 | 0.042258 |
| NM_030674 | SLC38A1 | 0.669886 | 0.020317 |
| NM_032926 | TCEAL3 | 0.666639 | 0.003534 |
| NM_014884 | SUGP2 | 0.654387 | 0.005024 |

**Supplementary Table 4:** DESeq2 results table showing significantly down regulated differentially expressed transcripts (DETs) in each condition. Each transcript is presented by both its unique RefSeq transcript ID and the related common gene ID/symbol, where possible. Transcripts not mapped to a specific gene ID are shown as 'None'. The log2FoldChange (L2FC) and adjusted P-value (padj) are noted. Transcripts are ordered by descending log2FoldChange.

| **DOWN REGULATED** | | | |
|---|---|---|---|
| **TRANSCRIPT ID** | **GENE SYMBOL** | **L2FC** | **PADJ** |
| **VEHICLE CONTROL VS 0.1 µM 3-HK** | | | |
| NM_134263 | SLC26A6 | -0.56924 | 0.007302 |
| NM_001278463 | DNM1L | -0.66023 | 0.035616 |
| NM_002693 | POLG | -0.68101 | 0.044335 |
| NR_073599 | None | -0.71123 | 0.013492 |
| NM_022911 | SLC26A6 | -0.74807 | 0.000371 |
| NM_001204747 | RFC1 | -0.76589 | 0.008655 |
| NM_012237 | SIRT2 | -0.8089 | 0.000242 |
| NM_199334 | THRA | -0.8894 | 0.003425 |
| NM_001278189 | TPM3 | -0.88945 | 0.001279 |
| NM_001273 | CHD4 | -1.80568 | 2.19E-05 |
| NR_110927 | SVIL-AS1 | -2.15756 | 0.004688 |
| NM_001813 | CENPE | -2.41438 | 0.000368 |
| NM_001100913 | PACS2 | -2.43777 | 0.02505 |
| **VEHICLE CONTROL VS 1 µM 3-HK** | | | |
| NM_001136472 | LITAF | -0.55277 | 0.008945 |
| NM_001024924 | EXOC1 | -0.57278 | 0.032852 |
| NM_001098616 | C1orf43 | -0.58819 | 0.029885 |
| NM_001015048 | BAG5 | -0.59052 | 0.048427 |
| NM_144669 | GLT1D1 | -0.63635 | 0.004494 |
| NM_012161 | FBXL5 | -0.65816 | 0.049676 |
| NM_031450 | C11orf68 | -0.74016 | 0.004113 |
| NM_001164380 | STAU2 | -0.74575 | 0.018124 |
| NM_001305155 | PPP2R3C | -0.78013 | 0.044034 |
| NM_001191005 | SRSF10 | -0.82913 | 0.024847 |
| NM_017958 | PLEKHB2 | -0.85581 | 0.026471 |
| NM_001668 | ARNT | -0.87793 | 0.011717 |
| NM_020831 | MRTFA | -0.92071 | 6.00E-08 |
| NM_013235 | DROSHA | -0.93972 | 0.037395 |
| NM_006444 | SMC2 | -0.99149 | 0.022238 |
| NM_001172677 | ZNF607 | -1.00123 | 0.006754 |
| NR_024111 | SBDSP1 | -1.00543 | 0.005025 |
| NM_001076785 | SLC7A6 | -1.07628 | 0.02513 |
| NM_005097 | LGI1 | -1.13666 | 0.004251 |
| NM_180699 | SNRNP35 | -1.22635 | 0.008974 |
| NM_001310332 | RNF31 | -1.28485 | 7.23E-05 |
| NM_001199355 | RPL17-C18orf32 | -1.30288 | 0.000391 |
| NM_001301819 | ZNF202 | -1.38181 | 0.000243 |
| NM_000093 | COL5A1 | -1.643 | 0.000137 |
| NM_024772 | ZMYM1 | -13.5351 | 1.63E-14 |
| **VEHICLE CONTROL VS 10 µM 3-HK** | | | |
| NR_024111 | SBDSP1 | -0.87275 | 0.0046 |
| NR_045116 | None | -0.88108 | 0.00944 |
| NM_001310332 | RNF31 | -1.25675 | 8.98E-07 |
| NR_024247 | PWWP3A | -1.35417 | 1.29E-08 |
| NM_031449 | ZMIZ2 | -1.5674 | 0.007655 |

| | | | |
|---|---|---|---|
| NR_125341 | DDX46 | -2.34708 | 0.000236 |
| NM_001206956 | CNTN4 | -2.58141 | 2.74E-06 |
| NM_001255986 | COLEC11 | -2.72058 | 0.012338 |
| **0.1 μM 3-HK VS 1 μM 3-HK** | | | |
| NM_001258217 | MIS12 | -0.55743 | 0.030625 |
| NM_032926 | TCEAL3 | -0.56567 | 0.011511 |
| NM_031450 | C11orf68 | -0.71664 | 0.00512 |
| NM_145276 | ZNF563 | -0.77244 | 0.006557 |
| NM_020831 | MRTFA | -0.83415 | 1.16E-06 |
| NM_001244813 | FOXP1 | -0.83693 | 0.032528 |
| NM_001267574 | EIF3C | -0.91665 | 0.014686 |
| NM_001199267 | DGKZ | -0.94507 | 0.030189 |
| NR_110801 | LOC100507002 | -0.95006 | 0.02239 |
| NM_032853 | PWWP3A | -0.95889 | 0.00884 |
| NM_004772 | NREP | -0.96608 | 0.004529 |
| NM_001164840 | LYRM4 | -0.98332 | 0.007484 |
| NM_001145468 | SPECC1L | -1.00764 | 0.04471 |
| NM_001310332 | RNF31 | -1.00779 | 0.002071 |
| NM_022107 | GPSM3 | -1.04824 | 0.006132 |
| NM_001128226 | DIS3 | -1.06 | 0.021007 |
| NM_001301856 | ELOVL5 | -1.06393 | 0.034458 |
| NM_001271951 | TPGS2 | -1.13824 | 0.002283 |
| NM_000093 | COL5A1 | -1.1624 | 0.006849 |
| NM_001172677 | ZNF607 | -1.1928 | 0.001049 |
| NM_001297553 | CHD4 | -1.20487 | 0.007721 |
| NM_003205 | TCF12 | -1.38335 | 0.015603 |
| NM_003646 | DGKZ | -1.47033 | 0.017771 |
| NM_001199746 | HOXD8 | -1.48397 | 0.001904 |
| NM_001287603 | ZBTB17 | -1.49822 | 0.034979 |
| NM_012472 | DNAAF11 | -1.57875 | 0.000538 |
| NM_001254749 | RGS5 | -1.59929 | 0.001229 |
| NM_001199355 | RPL17-C18orf32 | -1.6012 | 9.26E-06 |
| NM_001301819 | ZNF202 | -1.63701 | 1.02E-05 |
| NM_001014 | RPS10 | -1.73195 | 0.003079 |
| NM_197966 | BID | -1.95284 | 0.000233 |
| NM_001278174 | ZNF33A | -3.00301 | 0.017528 |
| NM_024772 | ZMYM1 | -13.5906 | 7.50E-15 |
| **0.1 μM 3-HK VS 10 μM 3-HK** | | | |
| NM_024307 | GDPD3 | -0.56107 | 0.002675 |
| NR_002450 | SNORD68 | -0.60962 | 0.037857 |
| NR_046325 | PCAT6 | -0.60994 | 0.016142 |
| NR_024247 | PWWP3A | -0.67704 | 0.01256 |
| NM_001105659 | LRRIQ3 | -0.7142 | 0.016433 |
| NM_001003788 | STRADA | -0.79853 | 0.041356 |
| NM_003250 | THRA | -0.83323 | 0.018595 |
| NR_038378 | LOC441242 | -0.92044 | 0.006438 |
| NM_003432 | ZNF131 | -0.92561 | 0.02602 |
| NM_002982 | CCL2 | -0.94799 | 0.000416 |
| NM_004772 | NREP | -0.95447 | 0.000619 |
| NM_001310332 | RNF31 | -0.9925 | 0.000154 |
| NM_015493 | KANK2 | -1.73523 | 0.008761 |
| NM_013274 | POLL | -1.74004 | 0.019582 |
| NR_130727 | LOC401357 | -1.80657 | 0.00055 |
| NM_003646 | DGKZ | -1.85346 | 0.000286 |

| | | | |
|---|---|---|---|
| NM_001254749 | RGS5 | -1.88052 | 1.91E-06 |
| NM_001256163 | BIRC2 | -2.19009 | 0.002988 |
| **1 µM 3-HK VS 10 µM 3-HK** | | | |
| NM_003099 | SNX1 | -1.27184 | 0.026831 |
| NR_024247 | PWWP3A | -1.36091 | 6.60E-06 |
| NR_130727 | LOC401357 | -1.54988 | 0.012869 |
| NM_001079877 | RASA4 | -2.67497 | 0.006277 |
| NM_001242701 | CAMTA1 | -2.83485 | 0.003476 |
| NM_174971 | ST3GAL3 | -3.21388 | 0.000369 |
| NR_024176 | MKNK1 | -3.45816 | 0.006483 |
| NM_001297778 | NMNAT1 | -3.81124 | 0.002374 |
| NM_001282660 | MRTFA | -3.89904 | 6.22E-13 |
| NM_001164824 | SMIM12 | -5.47745 | 0.00168 |

**Supplementary Figure 5:** Significant differentially expressed transcripts (DETs) per condition presented as Venn diagrams to highlight the number of overlapping DETs between experimental conditions. Data does not discriminate between up regulated or down regulated DETs. (A) Overlapping DETs for 0.1, 1 and 10 µM 3-HK vs vehicle control (0.02% DMSO). (B) Overlapping DETs for 0.1, 1 and 10 µM 3-HK compared to each other.

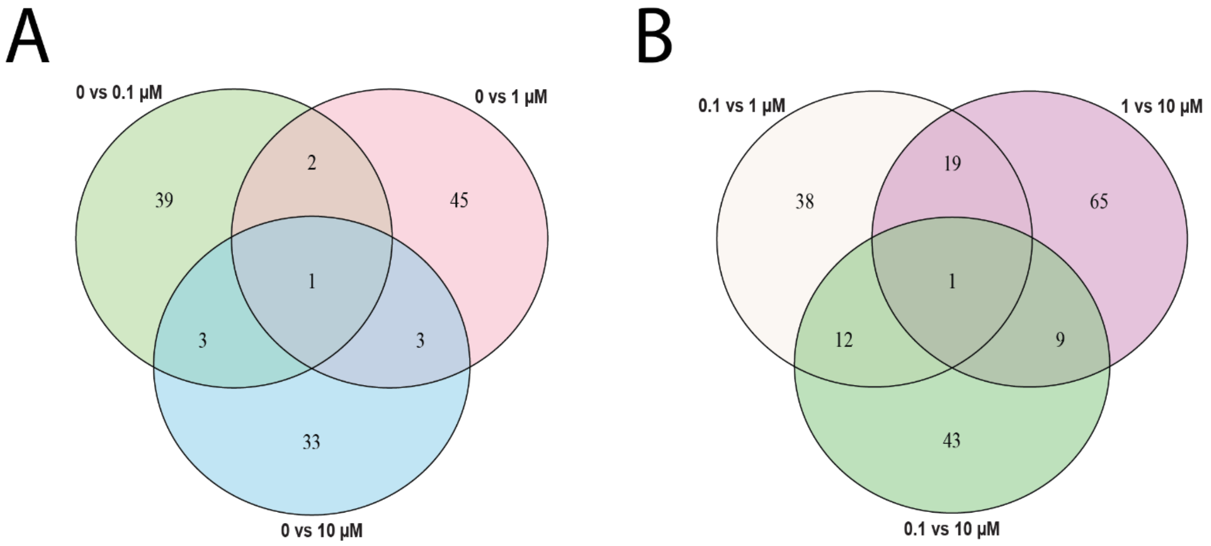

**Supplementary Table 5:** Table presenting the transcript and related gene names of significantly differentially expressed transcripts (DETs) which are up/down regulated in more than one experimental condition. Data does not discriminate between up regulated or down regulated DETs.

| TRANSCRIPT ID | GENE SYMBOL |
|---|---|
| **VEHICLE CONTROL vs 0.1 µM 3-HK VS VEHICLE CONTROL vs 1 µM 3-HK** | |
| NM_001166686 | PFKM |
| NM_003260 | TLE2 |
| **VEHICLE CONTROL vs 0.1 µM 3-HK VS VEHICLE CONTROL vs 10 µM 3-HK** | |
| NM_001282585 | MXRA8 |
| NM_032853 | PWWP3A |
| NM_001278393 | USP44 |
| **VEHICLE CONTROL vs 1 µM 3-HK VS VEHICLE CONTROL vs 10 µM 3-HK** | |
| NM_001310332 | RNF31 |
| NR_024111 | SBDSP1 |
| NM_001282568 | ZCCHC17 |
| **VEHICLE CONTROL vs 0.1 µM 3-HK VS VEHICLE CONTROL vs 1 µM 3-HK VS VEHICLE CONTROL vs 10 µM 3-HK** | |
| NM_001009812 | LBX2 |
| **0.1 µM 3-HK vs 1 µM 3-HK VS 0.1 µM 3-HK vs 10 µM 3-HK** | |
| NM_001205179 | ALKBH2 |
| NM_001813 | CENPE |
| NM_001273 & NM_001297553 | CHD4 |
| NM_001173482 | CRBN |
| NM_001105540 & NM_003646 & NM_001199267 | DGKZ |
| NM_004772 | NREP |
| NM_001303108 | PGAP4 |
| NM_001277335 | RASA4B |
| NM_001254749 | RGS5 |
| NM_001310332 | RNF31 |
| NM_012237 | SIRT2 |
| NM_001278189 | TPM3 |
| **0.1 µM 3-HK vs 1 µM 3-HK VS 1 µM 3-HK vs 10 µM 3-HK** | |
| NM_197966 | BID |
| NM_000093 | COL5A1 |
| NM_012472 | DNAAF11 |
| NM_001267574 | EIF3C |
| NM_022107 | GPSM3 |
| NM_001199746 | HOXD8 |
| NM_001013406 & NM_194454 | KRIT1 |
| NM_001282660 & NM_020831 | MRTFA |
| NM_001297778 | NMNAT1 |
| NM_001199355 | RPL17-C18orf32 |
| NM_001014 | RPS10 |
| NM_001164824 | SMIM12 |
| NM_148955 & NM_003099 | SNX1 |
| NM_174971 & NM_174963 | ST3GAL3 |
| NM_032926 | TCEAL3 |
| NM_024772 | ZMYM1 |
| NM_003455 & NM_001301819 | ZNF202 |
| NM_001278174 | ZNF33A |
| NM_001172677 | ZNF607 |
| **0.1 µM 3-HK vs 10 µM 3-HK VS 1 µM 3-HK vs 10 µM 3-HK** | |
| NM_006317 | BASP1 |

| | |
|---|---|
| NR_130727 | LOC401357 |
| NR_040013 | LOC644554 |
| NM_001285450 | PDXDC1 |
| NR_033836 | PIGA |
| NM_001256335 | PTGES2 |
| NM_003169 | SUPT5H |
| NM_001098493 & NM_001098496 | ZNF419 |
| NM_001172225 | ZNF540 |
| **0.1 µM 3-HK vs 1 µM 3-HK VS 0.1 µM 3-HK vs 10 µM 3-HK VS 1 µM 3-HK vs 10 µM 3-HK** | |
| NM_032853 & NR_024247 | PWWP3 |
